# Comparative Characterization of Bronchial and Nasal Mucus Reveals Key Determinants of Influenza A Virus Inhibition

**DOI:** 10.1101/2024.09.17.613498

**Authors:** Marie O. Pohl, Kalliopi Violaki, Lu Liu, Elisabeth Gaggioli, Irina Glas, Josephine von Kempis, Chia-wei Lin, Céline Terrettaz, Shannon C. David, Frank W. Charlton, Ghislain Motos, Nir Bluvshtein, Aline Schaub, Liviana K. Klein, Beiping Luo, Walter Hugentobler, Ulrich K. Krieger, Thomas Peter, Tamar Kohn, Athanasios Nenes, Silke Stertz

## Abstract

Differentiated primary human respiratory epithelial cells grown at air-liquid interface have become a widely used cell culture model of the human conducting airways. These cultures contain secretory cells such as goblet and club cells, which produce and secrete mucus. Here, we characterize the composition of mucus harvested from airway cultures of nasal and bronchial origin. We find that despite inter-donor variability, the salt, sugar, lipid, and protein content and composition are very similar between nasal and bronchial mucus. However, subtle differences in the abundance of individual components in nasal versus bronchial mucus can influence its antimicrobial properties: The ability of mucus to neutralize influenza A virus varies with the anatomical origin of the airway cultures and correlates with the abundance of triglycerides and sialylated glycoproteins and glycolipids.

**Importance:** Respiratory mucus plays an important role during the transmission and infection process of microbes in the human respiratory tract. In case of influenza A virus, mucus stabilizes virions in infectious respiratory particles and droplets but hampers virus particles before they reach the respiratory epithelium through its physicochemical properties and the presence of sialylated decoy receptors. However, it is thus far not well understood which components of mucus mediate protection and inhibition. Our study now provides a comprehensive analysis of bronchial and nasal mucus from primary human airway cultures that can be used as a resource for future experimental designs and interpretations.

## Introduction

Air-liquid interface (ALI) cultures of primary human airway epithelial cells have become a widely used tool to study the biology of airway epithelia *in vitro*. Cells from different anatomical sites in the respiratory tract for ALI cultures are usually derived from donors undergoing surgical procedures in the airways or, in the case of nasal airway epithelial cells, are obtained through minimally invasive techniques such as nasal brushings (1–4). Following proliferation on collagen-coated porous filter inserts in transwell plates, cell differentiation is induced by removing cell culture medium in the apical compartment. Differentiation into a pseudostratified epithelium takes approximately three weeks. ALI cultures strongly resemble human airway epithelia *in vivo* (4–6). In addition, the availability of airway cells derived from healthy and diseased individuals makes ALI cultures attractive for studying pathologies of the respiratory tract such as chronic obstructive pulmonary disease, cystic fibrosis, and asthma (7). Key features of human respiratory epithelium such as the presence of specialized cell types (e.g. ciliated cells, club cells, goblet cells, and basal cells), paracellular diffusion barriers formed by tight junctions, and an apical mucus lining are recapitulated in ALI cultures of airway cells, which make them suitable to study the interactions of respiratory pathogens with the human respiratory tract (7, 8).

One hallmark of fully differentiated ALI cultures of human airway epithelial cells is the production of mucus and airway fluids by secretory cells (primarily goblet and club cells) (1, 9). Airway mucus is a watery, viscous solution consisting of highly glycosylated mucins and a variety of other proteins, lipids, salts, sugars, as well as cellular debris (10, 11). Mucus lubricates the conducting airways and protects from stresses such as inhaled particles, toxins, and microbes (10). Mucins are the major macromolecular component of airway mucus. Membrane-tethered mucins (such as MUC1, MUC4, MUC16, and MUC20) are components of the periciliary layer (PCL) on the apical side of the airway epithelium. Secreted mucins (in particular MUC5B and MUC5AC) build a dense meshwork of polymers and form a viscous gel layer on top of the PCL (12) thereby allowing for mucociliary clearance of entrapped particles and cellular debris.

Airway mucus plays an important role in the defense against respiratory pathogens such as influenza A virus (IAV): Viral particles get entrapped in the viscous mucus layer and can be removed by mucociliary clearance before they reach the airway epithelia (13). Immobilization of virions by mucus can occur via non-covalent interactions with mucus glycoproteins (14), through size-exclusion during mucus penetration (15, 16), and by attachment to decoy receptors present in mucus. For IAV, it has been shown that interaction of the viral hemagglutinin (HA) with sialic acid receptors present on mucins inhibits productive infection of target cells (13, 17–19). As a result, the neutralization capacity of airway mucus against IAV can vary between virus strains and depends on the receptor-destroying activity of the viral neuraminidase (20). In addition, airway mucus contains a variety of antiviral proteins (e.g. components of the complement cascade, lactoferrin, immunoglobulins, and host defense peptides) (21–23), which reduce virus infectivity. Besides its antiviral activity, airway mucus and other respiratory fluids play a protective role during IAV transmission. Indeed, mucus and other matrices rich in proteins and other organics increase virion stability and preserve virus infectivity in droplets and aerosol particles in the air as well as on surfaces and fomites (24–31).

It has been demonstrated previously that apical secretions from tracheobronchial ALI cultures are similar to natural human respiratory secretions (21) making them an ideal model for studying IAV-mucus interactions. While differences based on anatomical origin as well as inter-donor variability have been described regarding barrier function, epithelial integrity, and cell composition of airway ALI cultures (3, 32, 33), a detailed characterization of mucus derived from these cultures is currently lacking. In this study, we report and compare the composition of apical secretions from primary human airway ALI cultures of bronchial and nasal origin in a comprehensive analysis of salt, lipid, protein, and glycan content. To address inter-donor variability, we characterize mucus samples of cultures derived from three donors per anatomical site. Given the predominant role of airway mucus in IAV transmission and infection, we further determine the neutralization capacity of bronchial and nasal mucus against IAV. Our results show that bronchial mucus exhibits a stronger neutralization capacity against IAV compared to nasal mucus. However, this effect is IAV strain-dependent and correlates with the abundance of α2,6-linked sialoglycans, proteins, and lipids, particularly triglycerides, in the mucus samples. In summary, we provide a detailed and comprehensive dataset on the composition of bronchial and nasal mucus from different donors that can help identify determinants of virus inhibition.

## Materials and Methods

### Cells and viruses

A549 (ATCC #CRM-CCL-185) and Manin-Derby canine kidney (MDCK) cells (ATCC #CRL-2936) were cultured at 37°C, 5% CO_2_, and >80% relative humidity in Dulbecco’s modified Eagle’s medium (DMEM, Gibco) supplemented with 10% heat-inactivated fetal calf serum (FCS, Gibco), 100 U/mL of penicillin, and 100 µg/mL of streptomycin (#15140–122, Gibco). Primary human epithelial cells were purchased from Epithelix (#EP51AB). Cells were derived from nasal (NEpC) or bronchial (BEpC) tissue. From each anatomical region, 3 donors were used: BEpC_AB051 (male, 53 years old, Caucasian, non-smoker), BEpC_AB079 (male, 62 years old, Hispanic, non-smoker), BEpc_AB0839 (male, 59 years old, Caucasian, non-smoker), NEpC_AB038 (male, 73 years old, Caucasian, non-smoker), NEpC_AB060 (male, 50 years old, Caucasian, non-smoker), and NEpC_AB063 (female, 41 years old, Caucasian, non-smoker). Cells were cultured in airway epithelium basal growth medium (#C-212601, PromoCell) supplemented with an airway growth medium supplement pack (#C-39160, PromoCell) and 10 µM Y-27632 (#1251, Tocris). For differentiation, transwell plates with 12-mm (#CLS3460, Corning) filter inserts were coated with collagen: a 0.5 mg/mL collagen (#C7774, Sigma-Aldrich) stock in 0.5 M acetic acid (#100063.1000, Merck) was diluted to 0.15 mg/mL in PBS prior to coating the filters. A total of 10^5^ BEpCs or NEpCs were seeded onto the coated transwell filters in a 1:1 mixture of airway epithelium basal growth medium and DMEM, which was supplemented with an airway growth medium supplement pack (Gray’s medium), and grown until confluence was reached. For differentiation at ALI, the medium was removed from the apical compartment, and Gray’s medium in the basal compartment was supplemented with 150 ng/mL retinoic acid (#R2625, Sigma-Aldrich). Cells were cultured at ALI for a minimum of 28 days prior to use. Medium in the basal compartment was refreshed every 2–3 days. The integrity of the epithelia was monitored by determining the transepithelial electrical resistance (TEER) weekly using an ERS-2 meter (Millicell). To measure TEER in representative wells, Gray’s medium is added to the apical compartment for the duration of the measurements.

TEER is normalized to the area of the transwell insert and given as area unit resistance (Ω*cm^2^). We consider cultures with a normalized TEER above 300 as intact.

The IAV strains A/Brisbane/59/2007 (H1N1) and A/Brisbane/10/2007 (H3N2) were grown in 10-day-old embryonated chicken eggs. Virus stocks were titered by standard plaque assay on MDCK cells.

### Harvesting of mucus from BEpC and NEpC ALI cultures

To remove the viscous mucus from the ALI cultures, 150 µL sterile ultrapure H_2_O was added to the apical compartment of the 12mm transwell and incubated for 10-15 min at 37°C. Dissolved mucus was removed from the cultures through carefully pipetting the supernatant from the apical compartment. Mucus from different wells was pooled per donor and stored at −80°C. Mucus harvest was performed at 4, 6, and 8 weeks of ALI culture. Mucus samples from different time points were thawed, pooled, and aliquoted for each donor separately and stored at −80°C until further use. It should be noted that our harvesting protocol may affect absolute concentrations of our measurements of mucus components.

### Quantification of IAV infectivity

IAV infectivity was quantified by plaque assay on MDCK cells. Briefly, cells were seeded in 12-well plates and grown to 90-100% confluency. Samples were diluted in series using PBSi (PBS for infection; PBS supplemented with 1% P/S, 0.02 mM Mg^2+^, 0.01 mM Ca^2+^, and 0.3% bovine serum albumin (#A1595, Sigma-Aldrich); pH ∼7.3). Cells were washed with PBS and infected with 100 μL inoculum and incubated for 1 h at 37°C with 5% CO_2_, with manual agitation every 10 min. The inoculum was removed, and cells were covered with an agar overlay (MEM supplemented with 0.5 μg/mL TPCK-trypsin, 0.01% DEAE-dextran, 0.11% sodium bicarbonate, and 0.7% Oxoid agar (#LP0028-500G, Thermo Fisher)). Cells were incubated for 72h at 37°C and fixed with 3.7% formaldehyde (#47608-1L-F, Sigma) in PBS. Cells were stained with a 0.2% crystal violet solution (#HT901-8FOZ, Sigma) in water and 10% methanol (#M-4000-15, Fisher Chemical). Plaques were enumerated to determine the virus titer in PFU/mL.

### Immunofluorescence

Differentiated ALI cultures of bronchial or nasal origin were fixed with 3.7% paraformaldehyde in PBS and permeabilized with PBS supplemented with 50 mM ammonium chloride (#254134; Sigma-Aldrich), 0.1% saponin (#47036, Sigma-Aldrich), and 2% BSA (#A7906; Sigma-Aldrich). A mouse anti-β-tubulin IV antibody (#ab11315; Abcam, United Kingdom), a mouse anti-MUC5AC antibody (#ab3649; Abcam), a rabbit anti-P63 antibody (#ab124762; Abcam), and a rat anti-uteroglobin antibody (#MAB4218; R&D Systems, USA) were used to stain ciliated, goblet, basal, and club cells, respectively. A rabbit anti-ZO-1 antibody (catalog no. 61-7300; Thermo Fisher Scientific) was used to stain tight junctions. As secondary antibodies, anti-mouse IgG Alexa488 (#A-11029), anti-rabbit IgG Alexa546 (#A-10040), and anti-rat IgG Alexa647 (#A-21247) antibodies were used (all from Thermo Fisher Scientific). Nuclei were stained with DAPI (#10236276001; Sigma-Aldrich). Filters were mounted using ProLong Gold Antifade Mountant (#P36930; Thermo Fisher Scientific), and z-stack images were acquired using a DMi8 microscope (Leica, Germany) and processed using the THUNDER Large Volume Computational Clearing algorithm (Leica). Maximum projection images of z-stacks were generated using LAS X (Leica) and ImageJ software.

### Microneutralization assay

A549 cells were seeded into clear (TPP) flat bottom 96-well plates to reach a confluent monolayer. Pre-warmed mucus (37°C) was diluted in Opti-MEM (#31985047, Thermo Fisher). The virus was thawed on ice and diluted in cold Opti-MEM to reach the desired concentrations. For each virus strain, a multiplicity of infection (MOI) was used that resulted in ∼80%–90% infection of the cell monolayer and a robust signal for quantification. Equal volumes of virus-mucus dilutions were mixed and incubated on ice for 1 hour. As positive controls, viruses were added to an equal volume of Opti-MEM. As a negative/mock control, Opti-MEM without the virus was used. Cells were washed with DPBS before 50 µL of the pre-incubated virus mixtures were added per well. Cells were then incubated for 1 hour at 37°C. DMEM supplemented with 100 U/mL penicillin, 100 µg/mL streptomycin, 0.3% BSA (#A7906, Sigma-Aldrich), 20 mM HEPES (#H7523, Sigma-Aldrich), and 0.1% FCS (p.i. DMEM) was added to the cells, which were incubated for another 6 hours at 37°C. Following incubation, cells were washed with DPBS, fixed and permeabilized (see section immunofluorescence). To stain IAV NP, a mouse monoclonal antibody (HB65, #H16-L10-4R5) diluted in CB was used. Alexa Fluor 488 donkey α mouse IgG (H+L) (#A-21202, Thermo Fisher Scientific) diluted in CB was used as the secondary antibody. The fluorescent signal was acquired using an IncuCyteS3 (Sartorius). The area of infected cells was determined using IncuCyte ZOOM 2018A software. First, the average area of infected cells from the mock-infected wells was subtracted from the area of infected cells from the infected wells. Negative values were set to 0. Next, the fold change relative to the average of the signal from the positive control wells was calculated. The infection levels in the positive control wells defined the “maximal infection.” To determine the mucus dilution at which 50% of the cells were still infected (IC_50_) in GraphPad Prism 9.2.0, the *x* values were log-transformed, and non-linear regression was used to fit a curve through the acquired data. The log-transformation required the *x* values of the positive controls to be set to a value different from 0, even if no mucus was added. Therefore, *x* values were artificially set to values 100-fold larger and smaller than the highest tested matrix dilution factor and the lowest tested antibody concentration, respectively. The curve was chosen to be top constrained at *y*, i.e., relative infection, equal to 1, and the baseline value was set to 0.

### Lipid Analysis

Lipids were extracted from mucus samples (25 µL) with 125 µL of isopropanol. Extracts were centrifuged, and the resulting supernatants were directly analyzed by LC-MS/MS as described by Medina et al. (2023) (34). Sample extracts were analyzed by HILIC-MS/MS, using a dual-column setup coupled to tandem mass spectrometry. Analysis was performed on a Vanquish™ Duo UHPLC System coupled to a TSQ Altis triple-stage quadrupole mass spectrometer (Thermo Scientific, San Jose, CA, United States) in positive and negative ionization modes. The chromatographic separation was carried out on an Acquity Premier BEH Amide column (1.7 µm, 100 mm × 2.1 mm I.D., Waters, Milford, MA, USA). A dual-column setup allowed for the re-equilibration of the first column while analyzing the sample on a second column, thus significantly reducing the total analysis time per sample. The mobile phase was composed of A = 10 mM ammonium acetate in Acetonitrile:H_2_O (95:5) (pH = 8.2) and B = 10 mM ammonium acetate in Acetonitrile:H_2_O (50:50) (pH = 7.4). The linear gradient elution from 0.1% to 20% B was applied for 2 min, from 20% to 80% B for 3 min, back to initial conditions (from 80% B down to 0.1%B) in the next 3 min, followed by 4 min of re-equilibration at initial chromatographic conditions (0.1% B). Following 6-minute separation gradient in positive ionization mode, the first column is switched offline for conditioning while the second column is switched inline for 6-minute separation in negative ionization mode, resulting in a 12-minute overall analysis time per sample. The flow rate was 600 µL/min, column temperature 45 °C, and sample injection volume 2 µL. Optimized HESI source parameters were set as follows: voltage 3500 V in positive mode and −2500 V in negative mode, Sheath Gas (Arb) =60, Aux Gas (Arb) =15, Sweep Gas (Arb) =1 and Ion Transfer Tube Temperature 380 °C. Nitrogen was used as the nebulizer and Argon as collision gas (1.5 mTor). Vaporizer Temperature was set to 350 °C. Optimized compound-dependent parameters were used for data acquisition in timed-Selected Reaction Monitoring (t-SRM) mode.

#### Data (Pre)Processing

Raw LC-MS/MS data was processed using the Trace Finder Thermo Scientific analysis software. The peak areas (or extracted ion chromatograms (EICs) for the monitored MRM transitions) were translated into concentrations based on single-point calibration with internal standard spike for complex lipids. Data quality assessment was performed using pooled quality control (QC) samples analyzed periodically throughout the entire batch (34).

### Analysis of ions with ion chromatography (IC)

Nasal and bronchial mucus (200 uL) were diluted in 2 mL of ultrapure water (Milli-Q system, 18 MΩ.cm). The diluted solution was filtered with a 13 mm syringe filter hydrophilic PTFE with 0.22 μm pore size. The filtered samples were injected in the instrument after the addition of chloroform (5 μl). The main anions (Cl^−^, NO ^−^, SO ^2−^, HPO ^2−^, C O ^2−^) were analyzed by ion chromatography (IC) after separation on a Dionex AS18 column (4 x 250 mm). The anions were determined with gradient elution at 1 mL min^−1^ with 23 mM KOH as eluent, and an ASRS-300 4 mm suppressor in auto suppression mode was used with applied current 90 mA. For the cations (Na^+^, NH_4_ ^+^, K^+^, Mg^++^, and Ca^++^), a CS12A-5μm (3x150 mm) column with a CSRS-300 4 mm suppressor was used. Separation was achieved under isocratic conditions with methanesulfonic acid (MSA) eluent (20 mM) and a flow rate of 0.5 mL min^−1^. The detection limit ranged from 1 to 5 ppb for the main anions and cations.

### Total Organics

Nasal and bronchial mucus were analyzed for organic carbon (OC), with the thermal–optical transmission method, using a carbon analyzer developed by Sunset Laboratory Inc., Oregon. A total of 5 μL of liquid sample were added onto 1.5 cm^2^ pre combusted quartz filters. The thermal method used in this study (EUSAAR2) was modified from the method developed by the National Institute for Occupational Safety and Health (NIOSH). The EUSAAR2 protocol was: step 1 in He, 200 °C for 120 s; step 2 in He 300 °C for 150 s; step 3 in He 450 °C for 180 s; step 4 in He 650 °C for 180 s. For steps 1–4 in He/O_2_, the conditions are 500 °C for 120 s, 550 °C for 120 s, 700 ° C for 70 s, and 850 °C for 80 s, respectively (58). The instrument was calibrated with sucrose solution and the detection limit was 0.2 μg cm^2^.

### Virus stability in droplets

#### Matrix preparation

Experiments were conducted in five matrices: nasal mucus, bronchial mucus, simulated lining fluid (SLF), H_2_O containing NaCl 1.9 g/L and H_2_O containing NaCl 3.2 g/L. SLF was prepared following the recipe adapted from Bicer (59) as detailed in Luo et al. (31) and freeze-dried according to the method described by Hassoun et al. (64). Briefly, SLF was made up from Hank’s Balanced Salt Solution (HBSS) without phenol red (#55037C, Sigma), lyophilized albumin from human serum (#A3782, Sigma), human transferrin (#T8158, Sigma), 1,2-dipalmitoyl-sn-glycero-3-phosphocholine (DPPC #P0673), 1,2-dipalmitoyl-sn-glycero-3-phospho-rac-(1-glycerol) ammonium salt (DPPG #42647), cholesterol (#C8667), L-ascorbic acid (#A5960), uric acid (#U0881), and glutathione (#PHR1359), (all purchased from Sigma-Aldrich). SLF powder was resuspended in milli-Q water prior to use.

#### Inactivation of IAV in 1-μl droplets

Inactivation experiments in droplets were performed as described in Schaub et al. (25). Experiments were performed under controlled conditions in an environmental chamber (#35532, Electro-Tech Systems) at 23 ± 2 °C and 40% RH ± 3 %. Virus stock was diluted 1:10 in ultrapure water to reduce medium contaminants and spiked 1:5 into the matrix of interest to achieve a starting titer in the experimental solution of 4*106 PFU/mL. For each condition, three 1 μL droplets were deposited into individual wells of a 96-well non-binding microplate (#655901, Greiner Bio-One). The last deposited set of droplets was immediately collected (t=0) and subsequent droplets were collected at t=30, t=60 and t=120 minutes post-deposition. Droplets were collected by resuspension in 300 μL of PBSi and agitation of the droplet, aliquoted and frozen at −20°C until infectivity scoring and genome copy (GC) quantification. For each matrix, a 1-μL sample was taken directly from the IAV-spiked matrix at the beginning and at the end of the experiment and diluted in 300 μL of PBSi as a control for viral decay in bulk solution.

Viral inactivation was quantified as log(N/N_0_), where N is the number of infectious viruses in a droplet, and N_0_ is the initial number of infectious viruses in the droplet (t=0). The values were corrected for physical virus losses due to attachment to the well plate (GC/GC_0_), which was determined from the fraction of genomic copies (GC) recovered from the droplet compared to the initial number of genomic copies in a 1 μL droplet at t=0 (GC_0_).

#### Quantification of IAV genome copies by droplet digital PCR

Viral RNA was extracted from 70 μL of collected sample using the QIAamp Viral RNA Mini extraction kit (#52906,Qiagen) according to the manufacturer’s instructions. Nucleic acids were recovered in 2x 40 μL (80 μL total) elution buffer and stored at −20°C until analysis. 12 μL reactions per sample were prepared containing 1X OneStep Advanced Probe Mastermix, 1X OneStep Advanced RT mix, GC enhancer, 0.8 μM Forward Primer (TGG AAT GGC TAA AGA CAA GAC CAA T), 0.8 μM Reverse Primer (AAA GCG TCT ACG CTG CAG TCC), 0.4 μM probe (5’ FAM-TTT GTK TTC ACG CTC ACC GTG CCC-BHQ-1 3’), 3 μL template and 3.18 μL H_2_O. Reactions were added to a QIAcuity Nanoplate 8.5k (#250021, Qiagen) and viral genome copies were quantified using the QIAcuity droplet digital PCR system under the following conditions: 40 min at 50°C and 2 min at 95°C for reverse transcription and RT inactivation, respectively, followed by 40 cycles of 5 sec at 95°C for denaturation and 30 sec at 60°C for annealing and extension. A no-template control of ultra-pure water was included in every run.

### BCA assay and proteomic analysis of bronchial and nasal mucus samples

#### BCA assay

Protein concentration of mucus samples was measured using the Pierce^TM^ BCA Protein Assay kit (#23227, Thermo Scientific) by following the instructions on the manufacturer’s protocol. Absorbance was measured with a 2140 EnVision multilabel plate reader (Perkin Elmer) and the obtained data was analyzed and interpolated using GraphPad Prism 10. Mucus samples of each donor were handed over to the Functional Genomics Center Zurich (UZH) for proteome identification and quantification. 20 μg of protein per donor were used in a label-based, fractionation approach with 12 final fractions for MS, to allow detection of low-abundant proteins.

#### Sample digestion and clean up

The protein concentration was estimated using the Lunatic UV/Vis polychromatic spectrophotometer (Unchained Labs). For each sample, 15 µg of protein were used. After addition of 20% SDS/tris-HCl to reach a final concentration of 4% SDS/Tris-HCl, the samples were treated with High Intensity Focused Ultrasound (HIFU) for 1 minute at an ultrasonic amplitude of 100% and boiled at 95°C for 10 minutes. The proteins were reduced with 5 mM TCEP (tris(2-carboxyethyl)phosphine) and alkylated with 15 mM chloroacetamide at 30°C for 30 min in the dark. Samples were processed using the single-pot solid-phase enhanced sample preparation (SP3). The SP3 protein purification, digest and peptide clean-up were performed using a KingFisher Flex System (Thermo Fisher Scientific) and Carboxylate-Modified Magnetic Particles (35) (GE Life Sciences; GE65152105050250, GE45152105050250). Bead conditioning was done with 3 washes with water at a concentration of 1 µg/µl. Samples were diluted with 100% ethanol to a final concentration of 60% ethanol. The beads, wash solutions (80% ethanol) and samples were loaded into 96 deep well-or micro-plates and transferred to the KingFisher. Collection of beads, protein binding to beads, washing of beads, protein digestion (overnight at 37°C with a trypsin:protein ratio of 1:50 in 50 mM triethylammoniumbicarbonat (TEAB)) and peptide elution were carried out on the robotic system. The digest solution and water elution were combined and dried to completeness.

#### TMT labeling and peptide fractionation

50 µg TMT 6-plex reagent (Thermo Fisher Scientific) was dissolved in 5 μl of anhydrous acetonitrile (Sigma-Aldrich) and added to 15 µg peptides in 15 µl of 50 mM TEAB, pH 8.5. The solution was gently mixed and incubated for 60 min at room temperature. The reaction was quenched by adding 1.2 µl of 5% hydroxylamine (Thermo Fisher Scientific). The combined TMT sample was created by mixing equal amounts of each TMT channel together. Labeled peptides were offline pre-fractionated using high pH reverse phase chromatography. Peptides were separated on an XBridge Peptide BEH C18 column (130Å, 3.5 µm, 1.0 mm X 250 mm, Waters) using a 72 min linear gradient from 5-40% acetonitrile/9 mM NH4HCO2. Every minute a new fraction was collected and concatenated into 12 final fractions.

#### LC-MS/MS analysis

Mass spectrometry analysis was performed on an Orbitrap Exploris 480 mass spectrometer (Thermo Fisher Scientific) equipped with a Digital PicoView source (New Objective) and coupled to an M-Class UPLC (Waters). Solvent composition at the two channels was 0.1% formic acid for channel A and 0.1% formic acid, 99.9% acetonitrile for channel B. Column temperature was 50°C. Peptides were loaded on a commercial nanoEase MZ Symmetry C18 Trap Column (100Å, 5 µm, 180 µm x 20 mm, Waters) connected to a nanoEase MZ C18 HSS T3 Column (100Å, 1.8 µm, 75 µm x 250 mm, Waters). Peptides were eluted at a flow rate of 300 nL/min. After a 3 min initial hold at 5% B, a gradient from 5 to 22% B in 80 min and 22 to 32% B in another 10 min was applied. The column was cleaned after the run by increasing to 95% B and holding 95% B for 10 min prior to re-establishing loading condition for another 10 minutes.

The mass spectrometer was operated in data-dependent mode (DDA) with a maximum cycle time of 3 s, funnel RF level at 40% and heated capillary temperature at 275 °C. Full-scan MS spectra (350−1500 m/z) were acquired at a resolution of 120000 at 200 m/z after accumulation to a target value of 3000000 or for a maximum injection time of 45 ms. Precursors with an intensity above 5000 were selected for MS/MS. Ions were isolated using a quadrupole mass filter with 0.7 m/z isolation window and fragmented by higher-energy collisional dissociation (HCD) using a normalized collision energy of 34%. HCD spectra were acquired at a resolution of 30000 and maximum injection time was set to Auto. The normalized automatic gain control (AGC) was set to 100%. Charge state screening was enabled such that singly, unassigned and charge states higher than five were rejected. Precursor masses previously selected for MS/MS measurement were excluded from further selection for 20 s, and the exclusion window was set at 10 ppm. The samples were acquired using internal lock mass calibration on m/z 371.1012 and 445.1200. The mass spectrometry proteomics data were handled using the local laboratory information management system (LIMS) (36).

#### Data analysis

The acquired shotgun MS data were processed for identification and quantification using Fragpipe 19.0 (Philosopher 4.8.1). Spectra were searched against a concatenated Uniprot human reference proteome and Uniprot bovine reference proteome (reviewed canonical version from 2023-10-17 concatenated to its reversed decoyed fasta database and common protein contaminants) using MSFragger 3.5 and Percolator. TMT modification on peptide N-termini and lysine side chains as well as carbamidomethylation of cysteine were set as fixed modification, while methionine oxidation was set as variable. Enzyme specificity was set to trypsin/P allowing a minimal peptide length of 7 amino acids and a maximum of two missed cleavages. Reporter ion intensities were extracted with 20 ppm integration tolerance. For peptide and protein quantification the co-isolation filter was set to 50%. The R package prolfqua (37) was used to analyze the differential expression and to determine group differences, confidence intervals, and false discovery rates for all quantifiable proteins. The protein lists were filtered with a threshold of 1 log2 FC and an FDR of 0.05%. The analysis was run on the local computing infrastructure (38).

#### Gene overrepresentation analysis

The top 30 abundant proteins from all six mucus samples were analyzed for enrichment with regard to gene ontology GO, cellular compartment, biological process, and molecular function using gProfiler (version e110_eg57_p18_4b54a898). To correct for multiple testing the Benjamini-Hochberg method was used and FDR cutoff was set to 0.05.

### N- and O-glycomic analysis

For N-glycomic profiling, harvested cells were extracted with lysis buffer containing 7 M urea, 2 M thiourea, 10 mM dithioerythreitol in 40 mM Tris buffer with 1% Protease inhibitor (Roche). The cell membranes were disrupted by High Intensity Focused Ultrasound with 10 times 10 s sonication with 16 amplitudes and 1 minute on ice in between, and subsequent shaking for 4 hours at cold room. The protein extracts were alkylated with 100 mM iodoacetamide in the dark for 4 hours at 37°C. Ice-cold trichloroacetic acid was added to a final 10% w/v concentration and left for one hour. After centrifugation at 20,000 g for 30 minutes at 4°C, precipitated sample pellets were washed twice with ice-cold acetone and then lyophilized. Dry protein pellets were redissolved in 50 mM ammonium bicarbonate buffer (pH 8.5), 250 unit of benzonase nuclease (Sigma-Aldrich) was added and incubated for 30 minutes at 37°C, following by trypsin digestion overnight. After deactivating the activity of trypsin, protein mixtures were further treated with PNGaseF (New England Biolab). The released glycans were cleaned up according to previous studies (66). In brief, the mixtures of glycans and peptides were loaded onto Sep-Pak C18 (Waters). N-glycans were eluted by 0.5% acetic acid and de- N-peptides were eluted by 20% isopropanol with 5% acetic acid and followed by 40% isopropanol with 5% acetic acid. N-glycans and de-N-peptides fractions were dried by SpeedVac.

To release O-glycans, retained peptide fraction from C18 Sep-Pak was submitted to alkaline-reductive elimination in 100 mM NaOH containing 1.0 M sodium borohydride at 45°C for 18 h. The reaction was stopped by addition of glacial acetic acid adding concentrated acetic acid dropwise on ice until the fizzing stops. The samples were loaded on Dowex 50W-X8 (50-100mesh) cation-exchange resin to remove residual peptides. and O-glycans were collected in 5% acetic acid. After evaporation of 5% acetic acid, boric acid was removed by 10% acetic acid in methanol to the sample and evaporating to dryness under the stream of nitrogen at RT. Before MALDI-MS analyses, the N- and O-glycan samples were permethylated using the sodium hydroxide/dimethyl sulfoxide slurry method, as described by Dell et al. (65). The samples were dissolved in 20 µL of acetonitrile. 1 µL sample mixed with 10 mg/mL 2,5- Dihydroxybenzoic acid (Bruker) in 70% Acetonitrile with 1mM sodium chloride was spotted on MALDI target plate and analyzed by Bruker RapiFlex^TM^ MALDI-TOF-TOF. Permethylated high mannose N-glycans and glycans from fetuin were used to calibrate the instrument prior to the measurement. The laser energy for each analysis was fixed and the data were accumulated from 10,000 shots. The data was analyzed by GlycoWorkbench (Ref: DOI 4310.1021/pr7008252) and inspected manually as described in the previous study (Ref: DOI 10.1038/s41423-024-01142-0). For relative quantification, the data was first deisotoped and the peak height was used for the calculation based on the following equation:

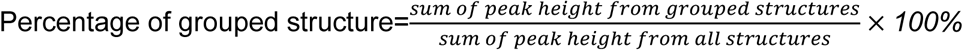

### SDS-PAGE and western blotting

Mucus samples were lysed in 5× Laemmli buffer (62.5 mM Tris-HCl pH 6.8, 25% glycerol, 2% SDS, 350 mM DTT, 0.01% Bromophenol Blue). Following treatment at 95°C for 5 min, mucus lysates (10 µg of proteins) were loaded onto a Bolt 4–12% Bis-Tris gradient gel (#NW04120BOX, Invitrogen). Proteins were separated by SDS-polyacrylamide gel electrophoresis (SDS-PAGE) followed by transfer to nitrocellulose membranes (#10600008, Amersham). Sialylated proteins were detected by western blotting using the following lectins: biotinylated Maackia Amurensis lectin I (MAL I, #B1315, vector laboratories, recognizes gal (β-1,4) glcNAc and gal (α-2,3) Neu5AC) and biotinylated Elderberry Bark lectin (EBL #B1305, vector laboratories, recognizes gal (α-2,6) Neu5AC). IRDye^®^ 800CW Streptavidin (#926-32230, Li-Cor) was used for detection with a LI-COR Odyssey Fc scanner.

## Results

### Mucus from bronchial and nasal epithelial cells differ in their inhibitory capacity against influenza A virus

Airway mucus not only varies in composition between individuals but also between the anatomical sites of production (39). However, a comprehensive and comparative dataset on mucus composition is currently lacking. We thus aimed to analyze the composition of mucus produced by primary human airway cultures derived from human bronchial or nasal tissue of a panel of donors. For each anatomical region, cells from three different donors of Caucasian or Hispanic origin were cultured in transwell plates. Bronchial epithelial cells (BEpC) and nasal epithelial cells (NEpC) were grown at air-liquid interface (ALI) to induce differentiation into a pseudostratified airway epithelium (**Fig 1A**). The integrity of the cultures indicated by their barrier function was monitored during differentiation by measuring the transepithelial electrical resistance (TEER) (32, 40, 41) (**Fig 1B, C**). Differentiation was confirmed through immunofluorescence staining of markers corresponding to ciliated cells (β-tubulin), goblet cells (MUC5AC), club cells (uteroglobin), and basal cells (p63) (**Fig 1D, E**). Mucus was collected from the apical, air-exposed side of the transwell culture at 4, 6, and 8 weeks of ALI culture through washes with ultrapure H_2_O, pooled, and stored at −80 °C until further use (**Fig 1A**). To test whether our bronchial and nasal mucus samples derived from different donors differ in their neutralization capacity against IAV we performed a microneutralization assay, in which IAV is pre-incubated with mucus before a monolayer of A549 cells is infected with the virus-mucus mixture. We chose two IAV strains, A/Brisbane/59/2007 (H1N1) and A/Brisbane/10/2007 (H3N2), which have been shown to differ in their sensitivity to neutralization by bronchial mucus (20). Bronchial mucus neutralized both virus strains with A/Brisbane/10/2007 (H3N2) exhibiting lower inhibitory concentration 50 (IC_50_) values compared to A/Brisbane/59/2007 (H1N1) confirming our previous findings (**Fig 2A-D**) (20). Interestingly, A/Brisbane/59/2007 (H1N1) was hardly neutralized by nasal mucus and IC_50_ values could be calculated only for mucus from two out of three donors (**Fig 2C, E**). When comparing the IC_50_ values of bronchial versus nasal mucus against A/Brisbane/59/2007 (H1N1), bronchial mucus had a significantly higher neutralization capacity (**Fig 2E**). The other virus strain, A/Brisbane/10/2007 (H3N2) was sensitive to neutralization by nasal mucus with IC_50_ values similar to bronchial mucus (**Fig 2F**). Our data indicate that bronchial and nasal mucus can neutralize IAV to different extents and that their neutralization capacity depends on the IAV strain.

**Figure 1.**
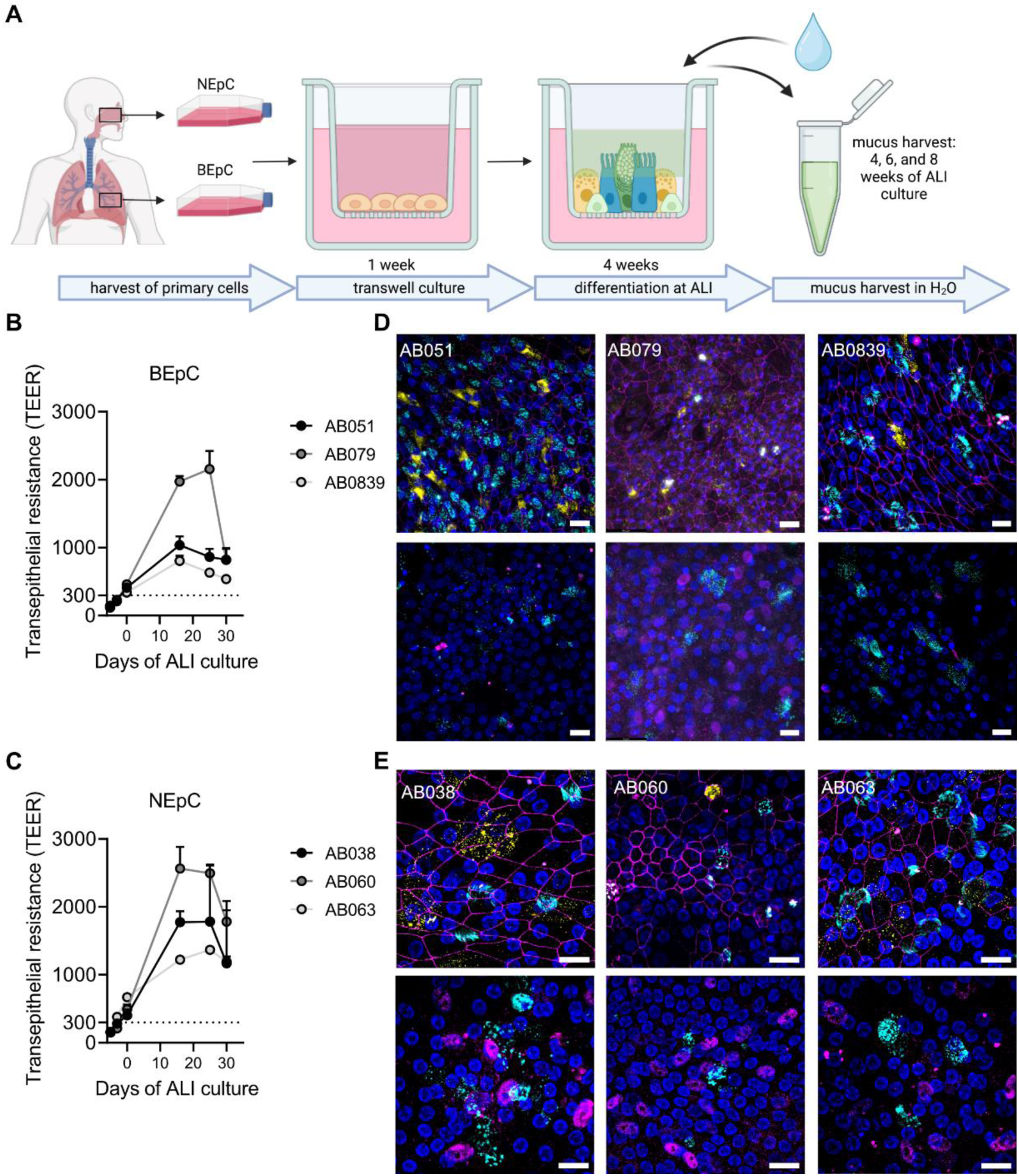
Differentiation of bronchial (BEpC) and nasal (NEpC) primary epithelial cells and mucus harvesting. **A** Schematic overview of primary cell differentiation and mucus harvesting. Following a four-week differentiation period at air-liquid interface (ALI), mucus was harvested in ultrapure H_2_O at 4, 6, and 8 weeks of ALI culture. The illustration was created at Biorender.com. **B-C** BEpC (B) and NEpC (C) from three different donors were differentiated and grown at an air-liquid interface (ALI) for 28 days. Transepithelial electrical resistance (TEER) was monitored at the time points indicated. Shown are means ± standard deviations from two independent wells. The dotted line indicates minimal TEER value for an intact cell monolayer. **D-E** Immunofluorescence staining of differentiated BEpC (D) and NEpC (E). Cells from three different donors of BEpC (AB079, AB051, AB0839) and NEpC (AB038, AB060, AB06) were fixed, permeabilized, and stained for the presence of ciliated cells (β-tubulin; cyan), club cells (uteroglobin, yellow), and tight junctions (ZO-1, magenta) shown in the upper panel. Goblet cells (Muc5AC; cyan), and basal cells (P63; magenta) are shown in the lower panel. Nuclei were stained with DAPI (blue). Z-stacks were transformed into maximum projection images. Scale bar represents 20 μm.

**Figure 2.**
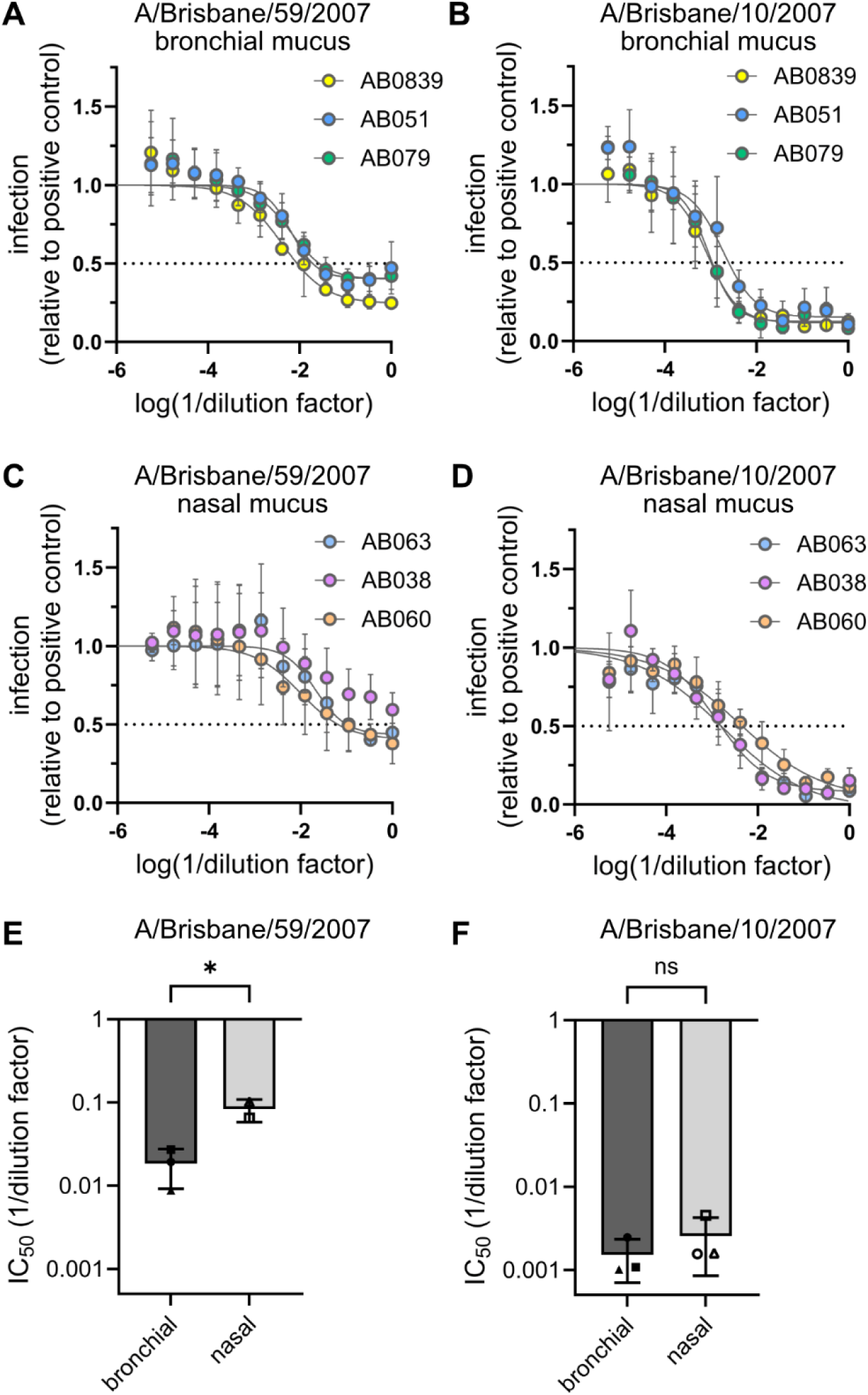
Bronchial mucus possesses a higher neutralization capacity than nasal mucus against A/Brisbane/59/2007 (H1N1) but not A/Brisbane/10/2007 (H3N2). **A-D** Neutralization capacity of bronchial (A, B) and nasal (C, D) mucus against A/Brisbane/59/2007 (H1N1, A, C) and A/Brisbane/10/2007 (H3N2, B, D). IAV was pre-incubated with serial dilutions of mucus for 1 hour on ice. A549 cells were then infected with the virus-matrix mixtures. For A/Brisbane/59/2007 (H1N1) a MOI of 5 was used while for A/Brisbane/10/2007 (H3N2) a MOI of 8 was used. At 7 hours post-infection, cells were fixed and stained for NP. The fluorescent signal relative to the positive control was quantified. Shown are means of three independent experiments. Error bars indicate standard deviation. Non-linear regression was used for curve fitting and to calculate the absolute IC_50_. **E, F** Absolute IC_50_ of bronchial and nasal mucus against A/Brisbane/59/2007 (H1N1, E) and A/Brisbane/10/2007 (H3N2, F) from three independent experiments. Significance was determined using a 2-way ANOVA and a Šídák’s multiple comparisons test. E: p=0.0006.

### Lipid, organics, and protein content but not salt concentrations correlate with the neutralization capacity of mucus against influenza A virus

In order to obtain a comprehensive and comparative dataset on mucus composition across donors and types of mucus that might help identify the determinants of neutralization capacity, we measured each mucus sample’s lipid, organic, protein, and salt content (**Table 1, Fig 3A, D, G, J)**. Notably, the lipid, organic, and protein content of bronchial mucus was higher than that of nasal mucus. One bronchial sample, AB0839, had a particularly high organics content of 1.34 C g/L which is two- to three-fold higher compared to other bronchial and nasal samples, respectively. In addition, this sample had the highest lipid (1.7 mg/L) and protein content (855.01 mg/L) (**Table 1**). Salt concentrations were similar between bronchial and nasal mucus and ranged between 1900 and 3714 mg/L (**Fig 3A**, **Table 1**). In a next step, we calculated the correlation for each mucus component with the mucus neutralization capacity (expressed as IC_50_, **Fig 2E-F**) against IAV (**Fig 3B-C, E-F, H-I, K-L**). The IC_50_ values of A/Brisbane/59/2007 (H1N1) correlated inversely with the lipid, organics, and protein content of the bronchial mucus samples but not well with those of the nasal mucus samples (**Fig 3E, H, K**). Furthermore, we could not detect a correlation between any of the measured mucus components in bronchial or nasal mucus and the IC_50_ values of A/Brisbane/10/2007 (H3N2) (**Fig 3C, F, I, L**). These results suggest that the increased neutralization capacity of bronchial mucus against A/Brisbane/59/2007 (H1N1) could be linked to the overall concentration of proteins and lipids in the bronchial samples which is two- to three-fold higher compared to the nasal mucus samples.

**Figure 3.**
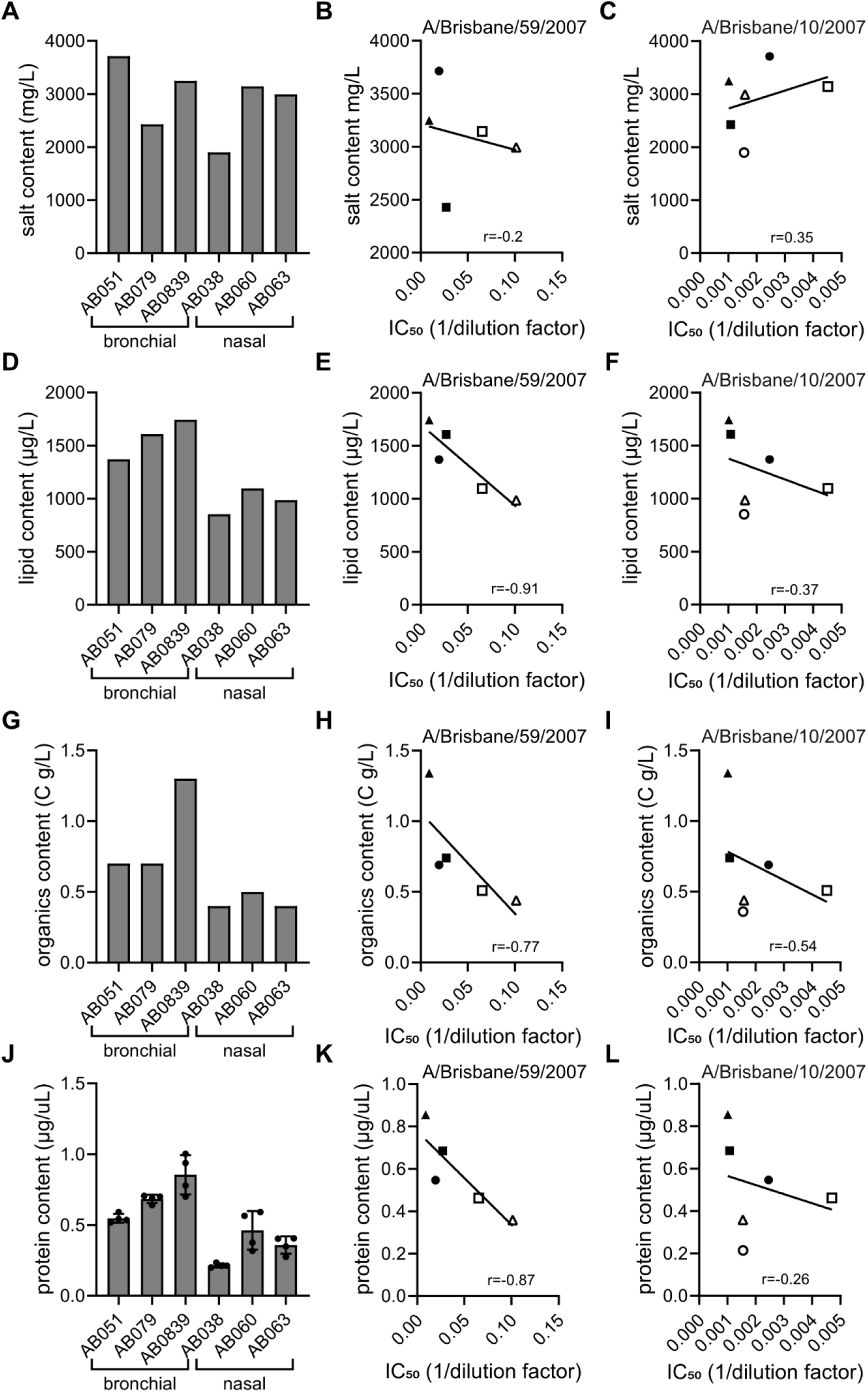
Correlation of mucus components with mucus neutralization capacity against A/Brisbane/59/2007 (H1N1) and A/Brisbane/10/2007 (H3N2). **A, D, G, J** Total salt, lipid, organics, and protein content from bronchial and nasal mucus samples. For G, shown are means of four independent measurements. Error bars indicate standard deviation. **B-C, E-F, H-I, K-L** Correlation between mucus lipid, organics, and protein content and IC_50_ against A/Brisbane/59/2007 (H1N1, middle panel) and A/Brisbane/10/2007 (H3N2, right panel), respectively. Shown is the linear correlation between the measurements of the mucus components from A, D, G, and J, and the IC_50_ values from Fig. 2E, F. r describes the direction and strength of the correlation. Bronchial mucus samples are depicted as closed symbols (AB051=circle, AB079=square, AB0839=triangle); nasal mucus samples are depicted as open symbols (AB038=circle, AB060=square, AB063=triangle).

**Table 1:** Composition of bronchial and nasal mucus in g/L.

| Sample | Salts | Lipids | Proteins | Organics* | Organics:salt ratio (g/g) |
| --- | --- | --- | --- | --- | --- |
| BepC-839 | 3.25±0.16 | 0.002 | 0.86± 0.14 | 1.34±0.09 | 0.41 |
| BepC-051 | 3.71±0.09 | 0.001 | 0.55± 0.03 | 0.69±0.05 | 0.18 |
| BepC-079 | 2.43±0.01 | 0.002 | 0.68± 0.03 | 0.74±0.06 | 0.3 |
| NepC-038 | 1.90±0.16 | 0.001 | 0.21± 0.01 | 0.36±0.04 | 0.19 |
| NepC-060 | 3.14±0.05 | 0.001 | 0.46± 0.14 | 0.51±0.05 | 0.16 |
| NepC-063 | 2.99±0.18 | 0.001 | 0.36± 0.06 | 0.44±0.04 | 0.15 |

\* Concentration of Organics is in C g L-1

### The organic content of bronchial and nasal mucus is sufficient to protect IAV from salt- mediated inactivation in 1 µL droplets

During transmission, mucins and other organic compounds of airway mucus stabilize IAV in infectious respiratory particles (43). In addition, a high organic:salt ratio provides a protective effect to IAV in drying respiratory droplets (25). We observed higher organics:salt ratios in bronchial versus nasal mucus (**Table 1**) which might indicate that respiratory droplets generated in the bronchi possess more protective properties compared to virus-containing droplets of nasal origin. Measurements of A/Brisbane/59/2007 (H1N1) stability in drying 1-µL droplets revealed a protective effect of airway mucus similar to simulated lining fluid (SLF) compared to NaCl solutions (**Suppl. Fig 1A**). However, no difference between bronchial and nasal mucus was observed in terms of virus stability. These data indicate that, outside the host, the complex composition and concentration of organics in mucus is sufficient to counteract the inactivating effect of salts over the time span considered.

### Salt analysis reveals no major difference between bronchial and nasal mucus

Variations in the composition of airway mucus can significantly alter its properties, for example through interactions of non-proteinaceous components with the mucin meshwork (42). We thus aimed to identify the abundant species of salts, lipids, proteins, and glycans in nasal versus bronchial mucus. When analyzing the salt composition of the mucus samples, we found that sodium and chloride were by far the most abundant, with concentrations ranging from 745 ± 49.57 to 1245 ± 3.6 and from 2023 ± 47.8 to 1070 ± 56.94 mg/mL, respectively. We also detected the presence of potassium, calcium, ammonium, and phosphate (**Suppl Fig 1B**). Overall, ion composition and concentrations were similar across most samples and none of the ions determined differed significantly between nasal and bronchial mucus. Interestingly, in mucus from one BEpC donor, AB051, increased concentrations of magnesium, sulfate, and nitrate ions were measured (**Suppl. Fig 1B**). However, these higher levels of the three ions did not correlate with differences in neutralization capacity.

### Lipidomic analyses reveal higher abundances of triglycerides in bronchial mucus

Lipidomics of the airway mucus samples revealed phosphatidylserines (PS), phosphatidylethanolamines (PE), and phosphatidylcholines (PC) as predominant lipid classes (**Fig 4A**). In particular, PS 18:0-18:1, PS 18:1-20:1, and PE 18:1-18:1 were highly abundant lipid species in bronchial mucus with average concentrations of 462.97±92.02, 122.58±2.77 and 89.55±9.64 µg/L, respectively (**Fig 4B**). The ratio between the different lipid species was similar across all samples (**Fig 4B**) but for most lipid classes, concentrations were higher in bronchial than in nasal mucus. To illustrate differential abundance, measurements from individual donors were grouped based on their anatomical origin, and fold changes of nasal mucus compared to bronchial mucus were calculated (**Fig. 4C**). Triglyceride (TG) lipids were particularly overrepresented within the fraction of lipids with lower abundances in nasal mucus. Other lipid species of differential abundance included individual sphingomyelin, hexosylceramide, and phosphatidylinositol (PI) species (**Fig 4C**). Notably, all differentially abundant lipid species were present at low concentrations (< 50µg/L) in airway mucus (**Fig 4B**). Interestingly, the TG and PI content correlated well with the neutralization capacity of mucus against A/Brisbane/59/2007 (**Suppl. Fig. 1 C-F**). These data indicate that lipid content differs qualitatively and quantitatively in bronchial and nasal mucus. Since lipids can affect mucus viscosity (43, 44, 60), lower levels of TGs in nasal mucus might lead to a less viscous and more fluid mucus layer than bronchial mucus.

**Figure 4.**
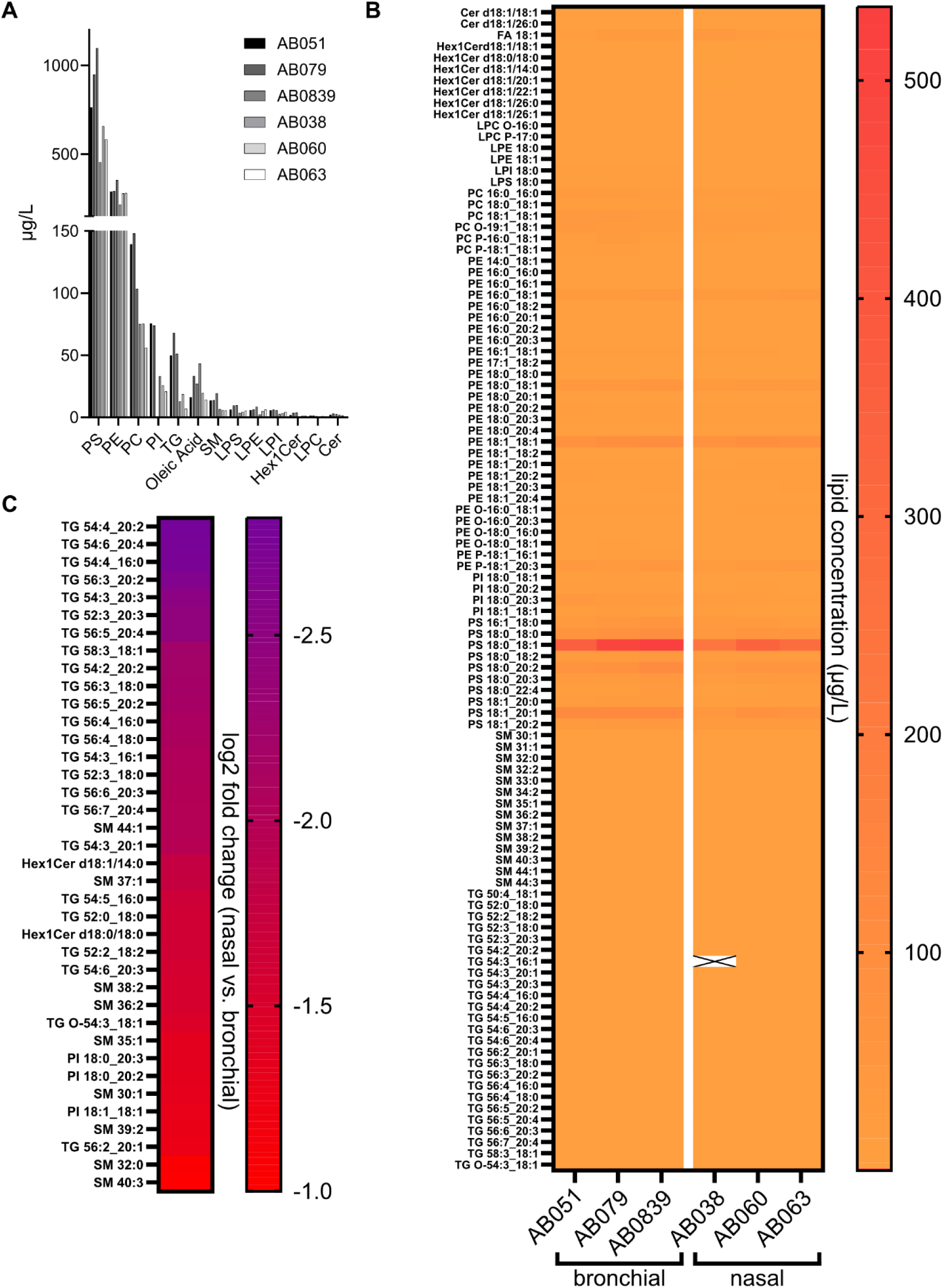
Lipid composition of bronchial and nasal mucus and differential abundance. **A, B** Abundance of lipid categories (A) and lipid species (B) in bronchial and nasal mucus samples. **C** Differential abundances of individual lipid species in nasal mucus versus bronchial mucus. Shown is the log2 fold change of lipid species in nasal mucus compared to bronchial mucus. Cut-off values for selection are a fold change of -2 and below and a p-value smaller than 0.05.

### The proteome of bronchial and nasal mucus reveals a high abundance of proteins linked to innate immune responses and secretory pathways

To investigate the protein composition of bronchial and nasal mucus, we used proteomic profiling of all mucus samples (**Fig 5, Suppl. Fig 2**). Hierarchical clustering of the relative abundances confirmed that samples from the same anatomical region are most similar and cluster together despite considerable inter-donor variability (**Suppl. Fig 2A**). In both bronchial and nasal mucus, highly abundant proteins were similar amongst the six donors regardless of the origin site, and are typically secreted proteins such as annexins, cathepsins, and uteroglobin (**Fig 5A**). In concordance with this observation, gene ontology analysis of each mucus sample’s 30 most abundant proteins confirmed the extracellular space as the most enriched cellular compartment (**Fig 5B**). Many of the highly abundant proteins possessed immunoregulatory functions or an association with the innate immune system: In particular, antimicrobial proteins such as components of the complement system, NGAL, and cystatin C were among the top 30 most abundant proteins (**Fig 5A**). Mucins are an integral component of respiratory mucus and are largely responsible for its gel-forming and barrier function. We found all major human airway mucins (MUC1, MUC4, MUC5AC, MUC5B, and MUC16) in the six mucus samples with high relative abundance values (**Fig 5C**) (12). Interestingly, mucin-1, a membrane-tethered mucin, rather than the secreted MUC5AB or MUC5B showed the highest relative abundance values (>20) of all detected mucins and was amongst the top 30 most abundant proteins (**Fig 5A, C**). Furthermore, we found that complement factor c3 was the most abundant protein in five out of six mucus samples (and the third most abundant protein in the remaining mucus sample). Within all mucus samples members of the complement cascade were present, many with high relative abundance values (**Fig 5D**). Overall, the protein composition of the six mucus samples was similar to what has been described for proteomic analyses of human airway secretions (21, 43). Next, we performed a differential expression analysis of nasal versus bronchial mucus, in which the abundances of the three individual donors per anatomical site were grouped, treated as independent replicates, and compared to each other. We found 25 proteins to be significantly higher expressed in nasal mucus while 41 proteins exhibited lower abundances (**Fig 5E, Suppl. Fig 2B**). The expression of uteroglobin, a component of airway fluids secreted by club cells, was reduced significantly in nasal mucus. This finding corresponds to club cells being primarily expressed in the distal airways (44). The majority of proteins expressed at reduced levels in nasal mucus were associated with metabolic processes (**Suppl. Fig 2C**). Amongst the lower abundant proteins in nasal mucus were a few proteins linked to innate immune defenses such as prohibitin-1/2 and lipopolysaccharide-binding protein (**Suppl. Fig 2B**). Mucus derived from nasal epithelial cells exhibited higher expression levels of proteins associated with metalloendopeptidase inhibitor activity and growth factor receptor binding (**Suppl. Fig 2D**). Interestingly, factors with antiviral functions such as tetherin and testican-2 were among the proteins with higher nasal abundances than bronchial mucus (**Suppl. Fig 2B**).

**Figure 5.**
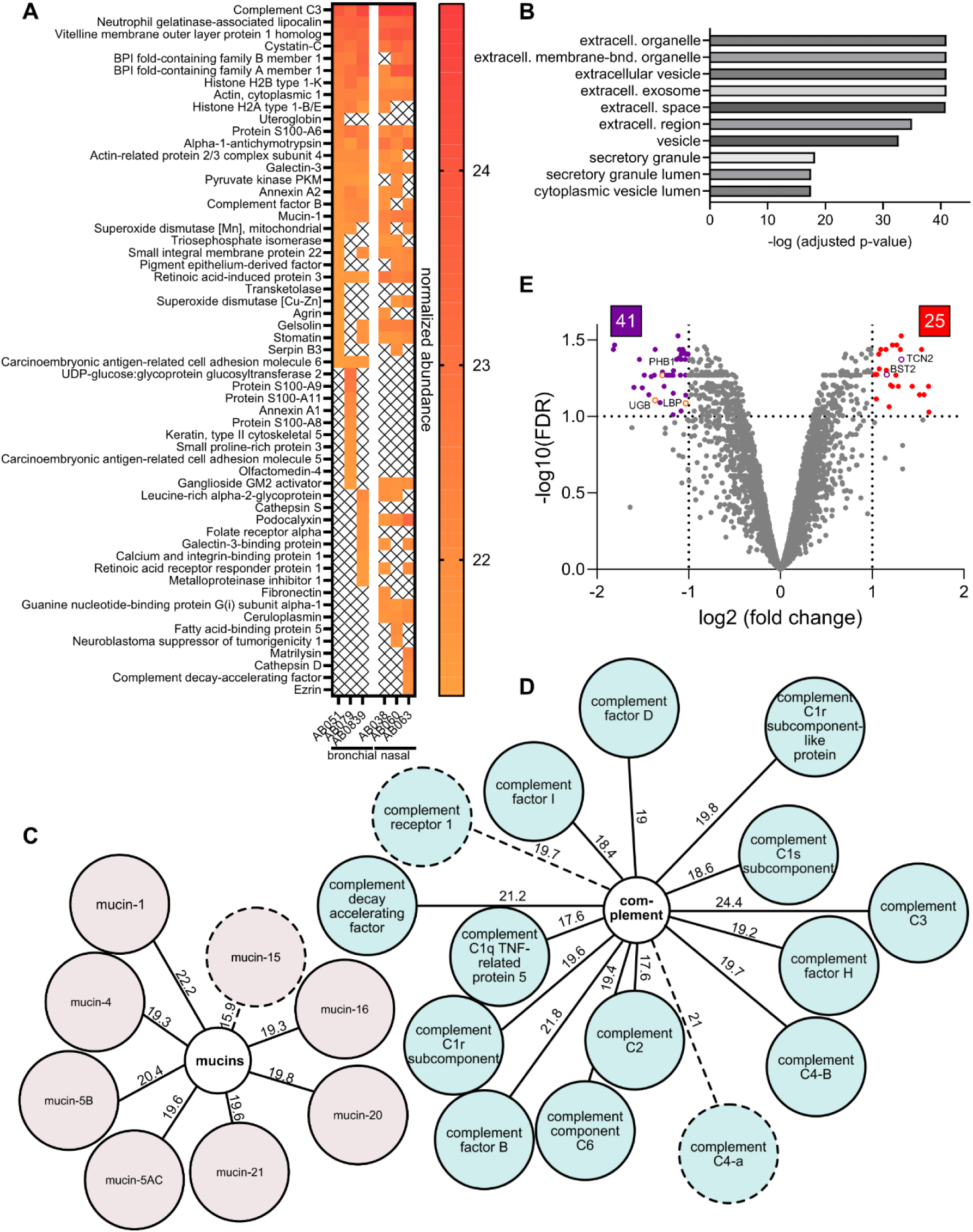
Proteomic profiling of bronchial and nasal mucus. **A** Top 30 protein abundance heatmap. Shown are the 30 most abundant proteins for each mucus sample ranked by the log 2 normalized abundance value. X indicates that a protein is not among the top 30 most highly abundant proteins of the respective mucus sample. **B** Gene Ontology analysis of the top 30 most abundant proteins shown in A. Analysis was performed with gene profiler and shown are the 10 most enriched GO:CC (cellular components) terms. **C, D** Mucins (C) and components of the complement system (D) detected in the mucus samples. Numbers indicate normalized protein abundance value. The dotted line indicates that only one peptide was identified for the respective protein. **E** Volcano plot showing the number of differentially abundant proteins in nasal mucus compared to bronchial mucus. Cut-off values (dotted lines) are a fold change of 2 and above and a false-discovery rate of 0.1 and below. Open circles depict examples of differentially abundant proteins with known pro- or antiviral function and uteroglobin.

### Bronchial mucus contains higher amounts of complex sialylated N-glycans with -2,6- linkage

After having analyzed salt, lipid, and protein composition we next determined whether the glycan repertoire differs between bronchial and nasal mucus. To this aim, we employed glycomics of the six mucus samples (**Fig. 6, Suppl. Fig 3-4, Suppl. Table 1-2**). Both bronchial and nasal mucus were found to contain various types of O-glycans and complex N-glycans (**Fig 6A-B, Suppl. Table 1-2, Suppl. Fig 3**). High amounts of N-glycans with core fucosylation were detected in addition to N-glycans with sialylation, sialyl Lewis structures, and polyLacNac with Lewis structures (**Figure 6C-F, Suppl. Fig 3, 4A-E, Suppl. Table 1**). Bronchial mucus was found to contain a significantly higher percentage of N-glycans with fucosylated core and terminal sialic acid (**Fig 6C**). In addition, we observed a trend that neutral N-glycans and O-glycans were more abundant in nasal mucus (**Fig 6D, Suppl. Fig 4I**). These glycan types correlated with a high and low bronchial and nasal mucus neutralization capacity against A/Brisbane/59/2007 (H1N1), respectively (**Fig. 6E, F, Suppl Fig 4J**). Furthermore, bronchial mucus contained higher overall amounts of sialic acid (**Fig 6G**), which correlated with the increased ability of bronchial mucus to neutralize IAV (**Fig. 6H**) supporting the hypothesis that sialic acid is the major driver of the neutralizing capacity of bronchial mucus. Given that human IAV strains have higher affinities to sialic acid bound to the underlying galactose via an α-2,6 glycosidic bond as compared to other linkages (62, 63), we next aimed to determine the levels of α-2,6-linked sialic acid. Lectin staining of Western Blots of the six mucus samples revealed that bronchial mucus contained more glycoproteins containing sialic acid with an α-2,6 linkage compared to nasal mucus (**Fig 6I, K**). The amounts of glycoproteins with α-2,3 linked sialic acid were similar between all samples (**Fig 6J, K**). In sum, in addition to differences in total sialic acid content bronchial mucus contains a selection of glycan-receptor types more suitable to neutralize human-adapted IAV strains.

**Figure 6.**
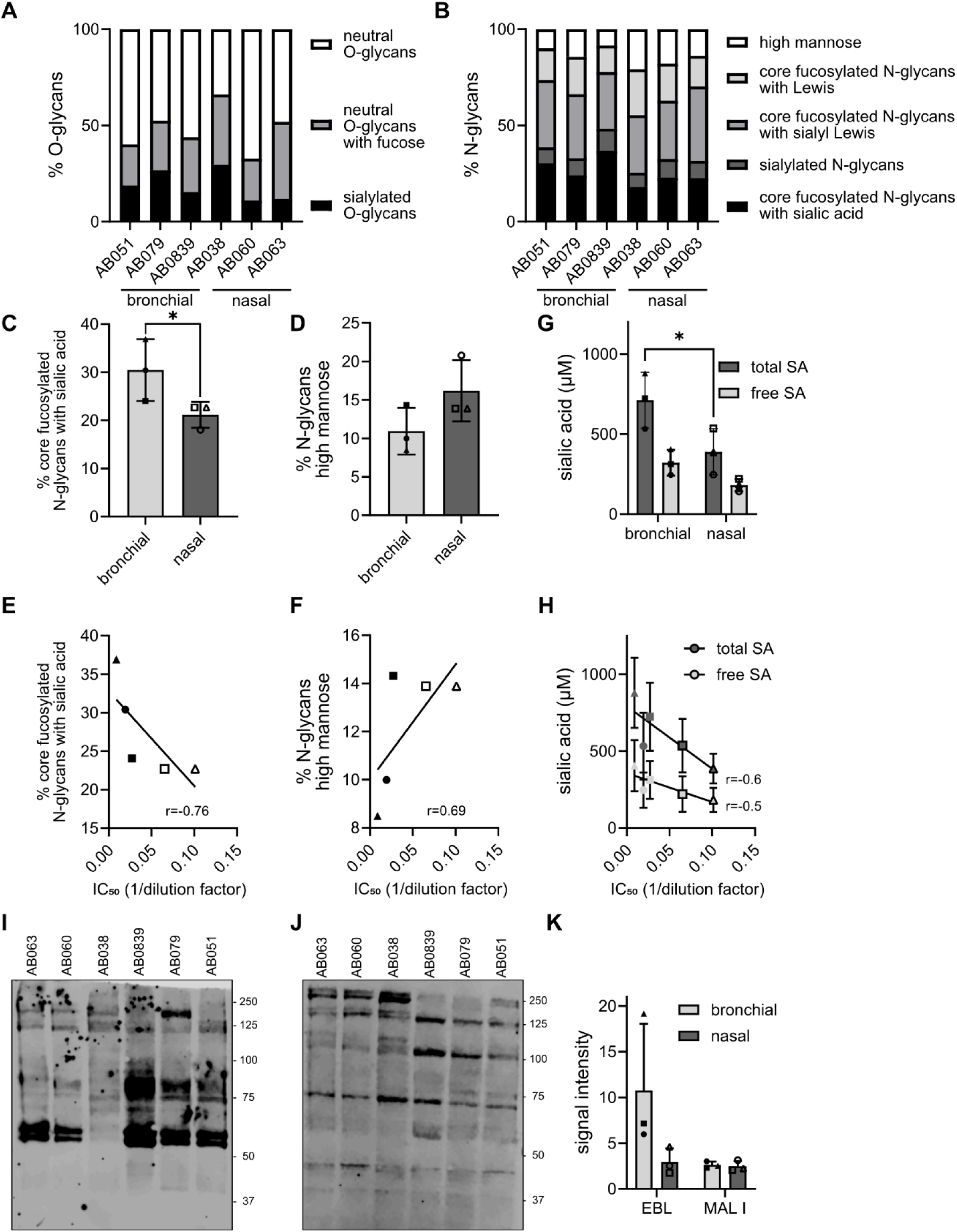
Glycan diversity of bronchial and nasal mucus. **A-B** Composition of O-glycans (A) and N-glycans (B) in bronchial and nasal mucus. **C** Percentage of sialylated N-glycans with fucosylated core. Shown are data from (A) analyzed by a 2-way ANOVA and a Šídák’s multiple comparisons test. p=0.026. **D** Percentage of neutral N-glycans. Shown are data from (A). **E-F** Correlation between N-glycan subtype abundance in mucus (C, D) and IC_50_ of mucus against A/Brisbane/59/2007. **G** Total and free sialic acid content of bronchial and nasal mucus. Shown are average values of three independent measurements. Significance was determined using a 2-way ANOVA and a Šídák’s multiple comparisons test. p= 0.0234. **H** Correlation between total and free sialic acid content (E) and IC_50_ of mucus against A/Brisbane/59/2007. r describes the direction and strength of the correlation. Bronchial mucus samples are depicted as closed symbols (AB051=circle, AB079=square, AB0839=triangle); nasal mucus samples are depicted as open symbols (AB038=circle, AB060=square, AB063=triangle). **I-K** Abundance of glycoproteins with α2,6 or α2,3 linked terminal sialic acid in bronchial and nasal mucus. Representative western blot of 10µg protein per mucus sample. α2,6 linked sialic acid containing glycoproteins were stained using Elderberry Bark lectin (EBL, I). α2,3 linked sialic acid containing glycoproteins were stained using Maackia Amurensis lectin I (MAL I, J). Signal intensity of three independent western blots was quantified (K).

## Discussion

A dual role has been suggested for airway fluids regarding IAV transmission: While virus infectivity is protected from damaging environmental factors in respiratory droplets and aerosols (25, 45), airway mucus represents a substantial barrier to the productive infection of the epithelia of the respiratory tract (13, 15, 18, 20, 46, 47). Our knowledge of the mechanisms underlying airway mucus’ pro- and antiviral activities during the different stages of IAV transmission is incomplete. With the advancement of 3D cell culture models of airway epithelia, human airway mucus can be obtained repeatedly in amounts large enough to address its role during IAV transmission in experimental setups. For the generation of human airway epithelial ALI cultures, cells are derived from human donors and are taken from different anatomical sites of the respiratory tract. Therefore, the composition and characteristics of mucus samples harvested from ALI cultures might vary substantially depending on the part of the respiratory tract and the individual from whom they were obtained. Indeed, survival of IAV in bronchial mucus derived from ALI cultures largely depended on individual cell culture donors (48). Yet, few studies have aimed at the characterization of mucus derived from airway epithelial ALI cultures (21, 43), and to our knowledge, none addressed donor or anatomical variability between different mucus samples. This study provides a detailed and comprehensive characterization of apical secretions from bronchial and nasal epithelial cells derived from a panel of donors.

We observed differences between the individual cultures in terms of epithelial integrity during differentiation of the airway cells into pseudostratified epithelia and in the cell composition of the cultures (**Fig1 B-E**), which is in line with reports from previous studies (32, 49). The transepithelial electrical resistance of all cultures was sufficiently high but declined after an early peak after fourteen days of ALI culture (**Fig 1B, C**) which is attributed to the development of a leaky barrier due to active secretions and apical-basal transport within the epithelium (50). In addition, all cultures contained specialized cell types typical for airway epithelium (**Fig 1D, E**). We therefore consider our cultures to be fully differentiated at the time of mucus harvest.

Substantial variability in mucus composition has been reported for bronchial mucus obtained from epithelial ALI cultures from different donors (48). Our proteomic profiling of the bronchial and nasal mucus samples reveals that the degree of variability in protein composition between samples from different anatomical sites is larger than the inter-donor variability (**Suppl Fig 2A**). Thus, bronchial and nasal mucus can be distinguished based on protein composition. Nevertheless, when looking at high-abundant proteins, all mucus samples were very similar regardless of their anatomical origin (**Fig 5A**). Proteins with functions linked to the extracellular space and innate immune responses were enriched in all samples (**Fig 5A, B**), which matches earlier characterizations of apical secretions from bronchial epithelial cultures grown at ALI and *ex vivo* human airway fluids (21, 43). Components of the complement system were strongly enriched in both bronchial and nasal mucus, with complement factor C3 being the most abundant protein of our analysis (**Fig 5D**). The importance of the complement system in innate defenses of the respiratory mucosa is well established (51). Besides components of the complement cascade, our data confirm the presence of a myriad of other antimicrobial and immune-regulatory factor as well as mucin-interacting proteins (**Fig 5A)** that have previously been linked to the innate defense capacity of airway mucus (52). Remarkably, MUC1 was the only mucin amongst the thirty most highly expressed proteins. Based on our harvesting protocol which enriches for soluble mucus components and analyses of sputum samples (53) we would have expected to find higher abundances of secreted mucins (such as MUC5B and MUC5AC) rather than membrane-tethered mucins (such as MUC1 and MUC4). While we identified eight mucins in our mucus samples **(Fig 5C**), unlike other proteomic analyses of airway secretions (11, 54), their abundances are likely underrepresented due to their large sizes and methodological limitations. The technical difficulties in detecting mucins using standard MS protocols have been identified before (21). It should be noted that mucus from ALI cultures differs from *in vivo* secretions due to the lack of submucosal glands, secretions from migratory cells, and “contaminations” by airway fluids from other anatomical regions and saliva. Indeed, certain highly abundant components of airway fluids, such as immunoglobulins, were absent from ALI culture-derived mucus (**Fig 5**). Nevertheless, our data show that mucus harvested from 3D airway epithelial cultures are suitable representatives of *in vivo* secretions and recapitulate anatomical site-specific differences in protein composition.

It is well-established that airway mucus reduces IAV infectivity (13, 17). Previously, we have observed that the susceptibility of IAV to neutralization by bronchial mucus varies, depending on the IAV strain used in a microneutralization assay (20). In this study, we tested the ability of bronchial and nasal mucus to neutralize a mucus-sensitive strain, A/Brisbane/10/2007 (H3N2), and a mucus-resistant strain, A/Brisbane/59/2007 (H1N1) and were able to recapitulate our earlier findings: for bronchial and nasal mucus, A/Brisbane/10/2007 (H3N2) was approximately one order of magnitude more sensitive to neutralization, as indicated by IC_50_. This implies that the susceptibility between IAV strains or isolates to inhibition by airway mucus is independent of its anatomical origin. Even though variations in mucus compositions will likely not alter the virus’ general sensitivity to neutralization by airway mucus we found that the mucus-resistant IAV strain, A/Brisbane/59/2007 (H1N1), was inhibited more strongly by bronchial mucus compared to the nasal samples. Thus, qualitative differences between bronchial and nasal mucus affect the antiviral properties of airway mucus against certain IAV strains. Of note, inter-donor variability did not play a role in the observed difference in neutralization potential as the IC_50_ for the individual mucus samples of the same anatomical origin were of similar magnitude (**Fig 2E, F**).

Further characterization of the mucus samples found indeed differences in composition between bronchial and nasal mucus: Bronchial mucus contained significantly more proteins and lipids compared to nasal mucus (**Fig 3 and 5**) and the total protein and lipid content correlated inversely with the IC_50_ against A/Brisbane/59/2007 (H1N1). When characterizing the lipid species present in our mucus samples, we found, similar to previous reports (61), phospholipids such as PS, PE, and PC to be highly abundant. The various lipids present in mucus contribute differently to the physicochemical properties. Surface-active phospholipids, improve the wettability of mucus, whereas neutral lipids such as glycerolipids e.g., triglycerides (TG) and glycosphingolipids contribute to the viscosity of mucus (60). TG were, amongst other lipid species, significantly enriched in bronchial mucus (**Fig 4C**). Lower levels of TGs in nasal mucus might lead to a less viscous and more fluid mucus layer compared to bronchial mucus. This difference can impact the mucus’s ability to trap and transport particulates and pathogens. Lipid profiles in airway fluids have been linked previously to respiratory infections: Humes and colleagues characterized sputum samples from individuals with respiratory infections and found that increased levels of TG and diacylglycerols (DG) were associated with IAV infections (55). Their study found an association with increased TG and DG levels for IAV infections with subtype H3 and not with subtype H1. Therefore, while high TG concentrations might be favorable for H3 infections, H1 infectivity might be negatively affected through mucus enriched in DG, as in our microneutralization assay (**Fig 2**).

We observed a trend towards a higher organics:salt ratio in bronchial mucus which might affect the survival of IAV outside the host in respiratory droplets (25). Interestingly, both nasal and bronchial mucus preserved virus infectivity in droplets unlike organic-free saline solution at the same salt concentration as the mucus (**Suppl. Fig 1C**). Likely, respiratory fluids, regardless of their anatomical origin, contain sufficiently high organics concentrations to protect IAV in infectious respiratory particles.

It is tempting to speculate that individual proteinaceous components are responsible for the observed variability in the neutralization capacity of airway mucus from different anatomical origins. Our differential expression analysis of the proteomic mucus profiles found proteins with immune-regulatory and antiviral activity to vary in abundance between bronchial and nasal mucus: Prohibitin-1 (PHB1) is higher expressed in bronchial mucus while testican-2 and bone marrow stromal antigen 2 (BST-2) exhibits increased abundance in nasal mucus (**Fig 5E, Suppl. Fig 2B**). The expression profiles of these respective pro- and antiviral proteins do not correspond with our neutralization data against A/Brisbane/59/2007 (H1N1) (**Fig 2**). However, higher overall protein and lipid levels in bronchial mucus compared to the nasal samples (**Fig 3**) likely translates into an increased presence of available decoy receptors in bronchial mucus and might outweigh the increased expression of individual proteins with confirmed antiviral activity against IAV. Indeed, the overall abundance of sialic acid, particularly sialic acid with an α-2,6 type linkage, was higher in bronchial mucus than in nasal mucus, supporting this hypothesis (**Fig 6I-K**).

To our surprise, we found that the mucus-sensitive IAV strain, A/Brisbane/10/2007 was neutralized equally well by bronchial and nasal mucus. This observation raises the question of whether there is a yet unknown, IAV strain-dependent factor that determines the virus sensitivity to neutralization by mucus with differences in composition (e.g. derived from different anatomical sites). We showed previously that the neuraminidase activity of an IAV strain correlates with its resistance to neutralization by bronchial mucus (20). Therefore, we can speculate that IAV strains with low neuraminidase activity, such as A/Brisbane/10/2007, are readily neutralized by mucus’s large abundance of decoy receptors. This idea is further supported by the high abundance of a mixture of complex sialoglycans in airway mucus to which H3 IAV strains have been shown to bind with high avidity (56, 57). On the other hand, IAV strains with high neuraminidase activity might be able to counteract immobilization through decoy receptors up to a certain threshold. If that threshold is met by the receptor-destroying capacity of an IAV strain, variations in the amounts of well-binding sialoglycan decoys might influence a strain’s sensitivity to neutralization.

In summary, this study provides a detailed and comprehensive characterization of mucus from human airway epithelial cells of bronchial and nasal origin, which can help identify determinants of virus inhibition or protection of infectivity in studies on IAV transmission.

## Figure Legends

**Suppl. Figure 1.**
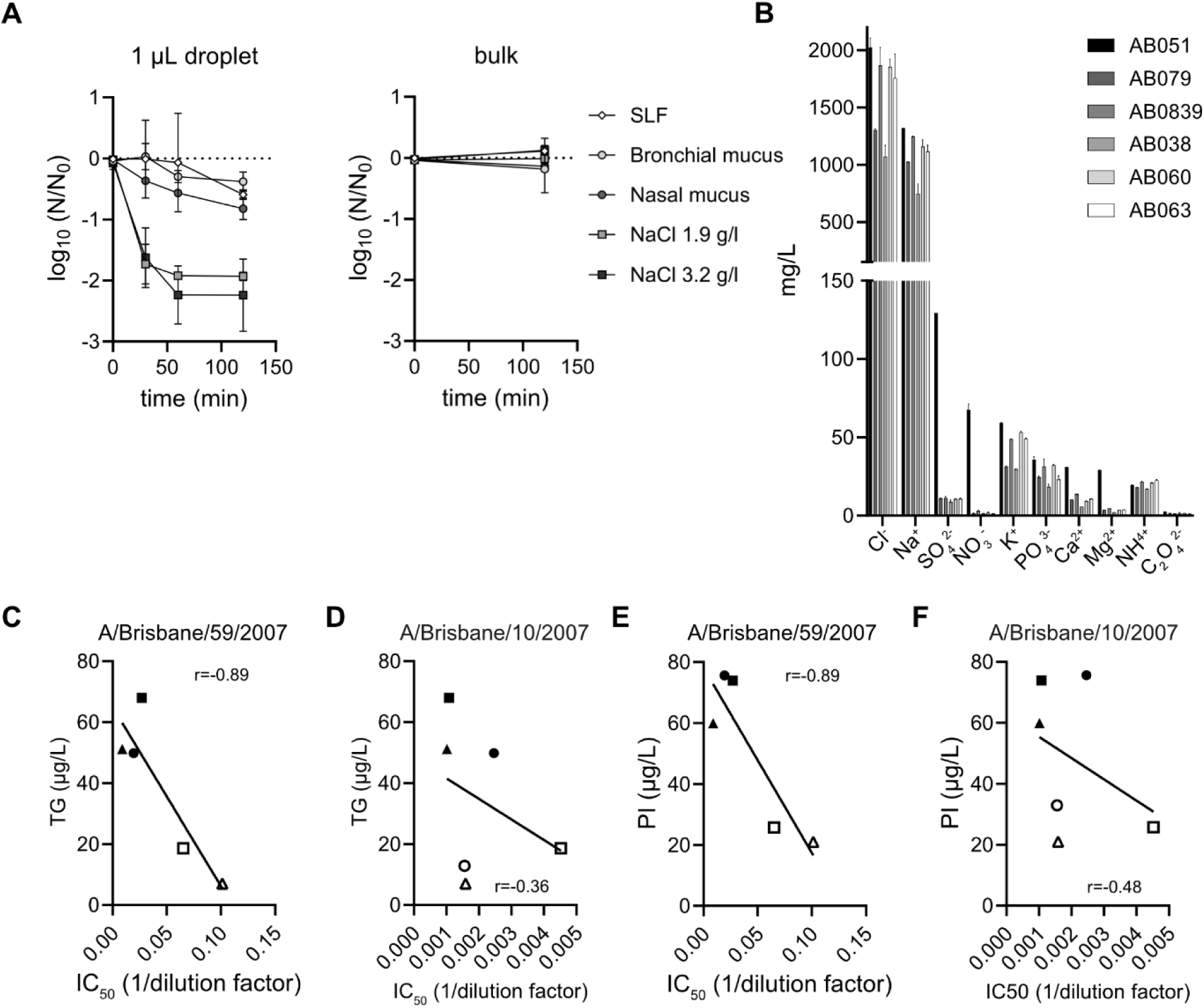
**A** Virus stability in 1-µL droplets. A/Brisbane/59/2007 diluted in saline, SLF, bronchial (AB0839) or nasal (AB038) mucus was incubated for indicated time points at 40% RH and 25°C. Shown is virus decay in droplets and bulk over time relative to 0 min as log_10_ N/N_0_. **B** Abundance of different salts in the individual mucus samples. **C-D** Correlation between TG content and IC_50_ of mucus against A/Brisbane/59/2007 (C) and A/Brisbane/10/2007 (D). **E-F** Correlation between PI content and IC_50_ of mucus against A/Brisbane/59/2007 (E) and A(Brisbane/10/2007 (F). r describes the direction and strength of the correlation. Bronchial mucus samples are depicted as closed symbols (AB051=circle, AB079=square, AB0839=triangle); nasal mucus samples are depicted as open symbols (AB038=circle, AB060=square, AB063=triangle).

**Suppl. Figure 2.**
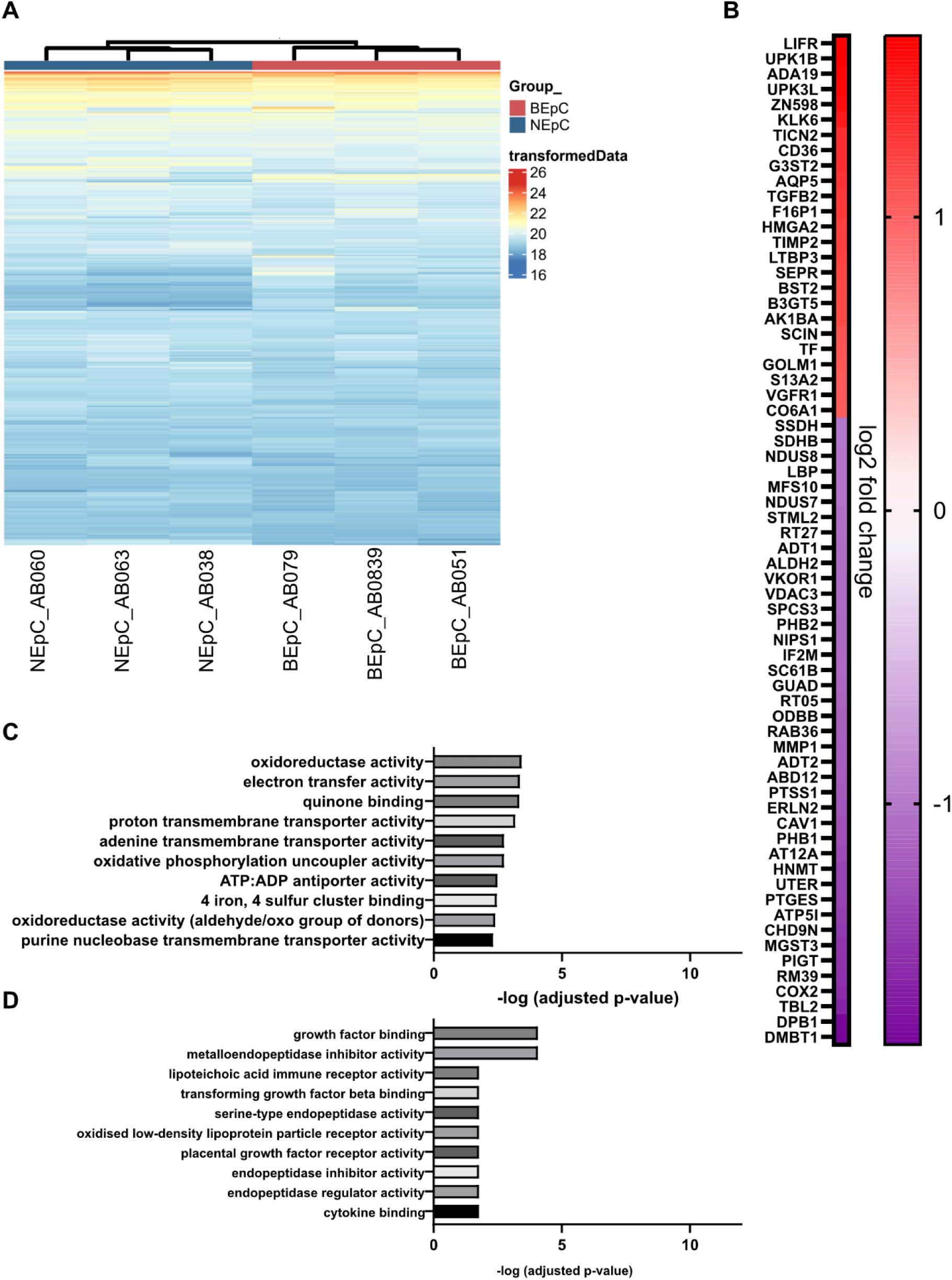
**A** Protein abundance heatmap. Rows indicate proteins; columns indicate mucus samples. Shown is the normalized protein abundance value of the top 1000 abundant proteins. Co-clustering (hierarchical complete linkage, euclidean distance) of samples and proteins was applied. **B** Heat map showing the differentially expressed proteins (gene ID) in nasal mucus compared to bronchial mucus. Proteins are ranked based on the log 2-fold change. **C, D** Gene Ontology analysis of the differentially expressed proteins in nasal mucus versus bronchial mucus (B). Analysis was performed with gene profiler and shown are the 10 most enriched GO:MF (molecular function) terms for proteins of lower (C) and higher (D) abundance in nasal mucus.

**Suppl. Figure 3.**
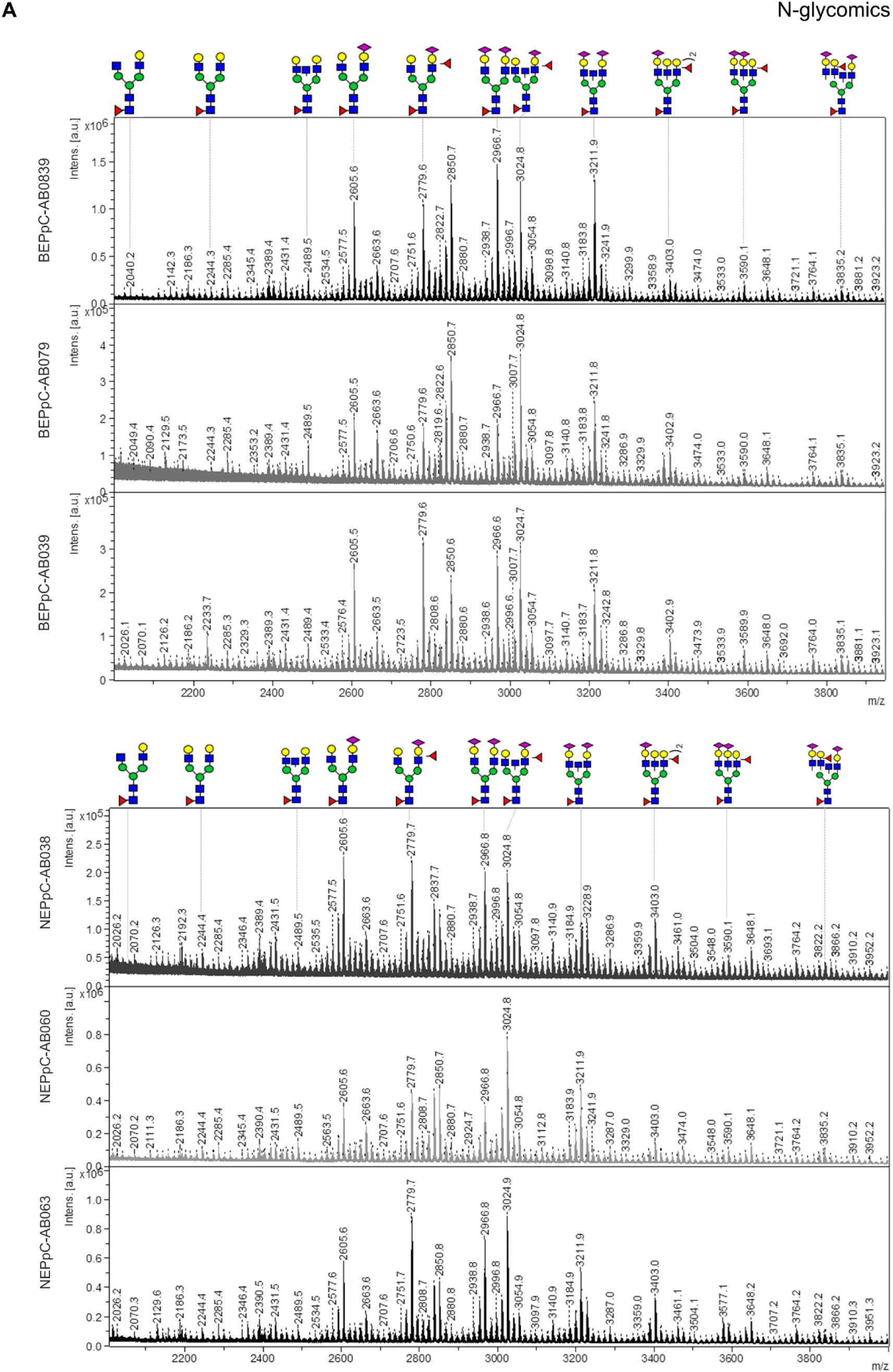
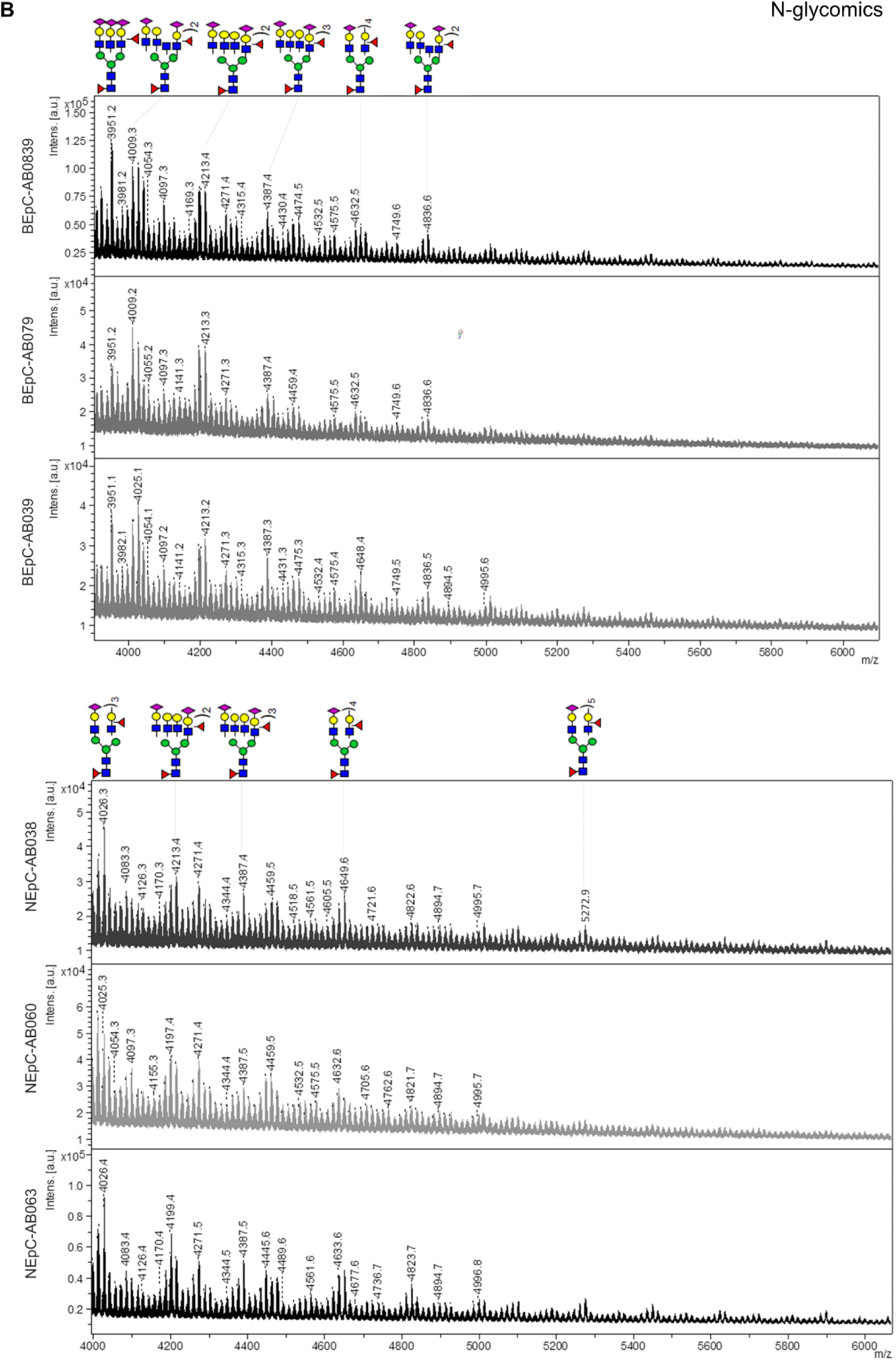
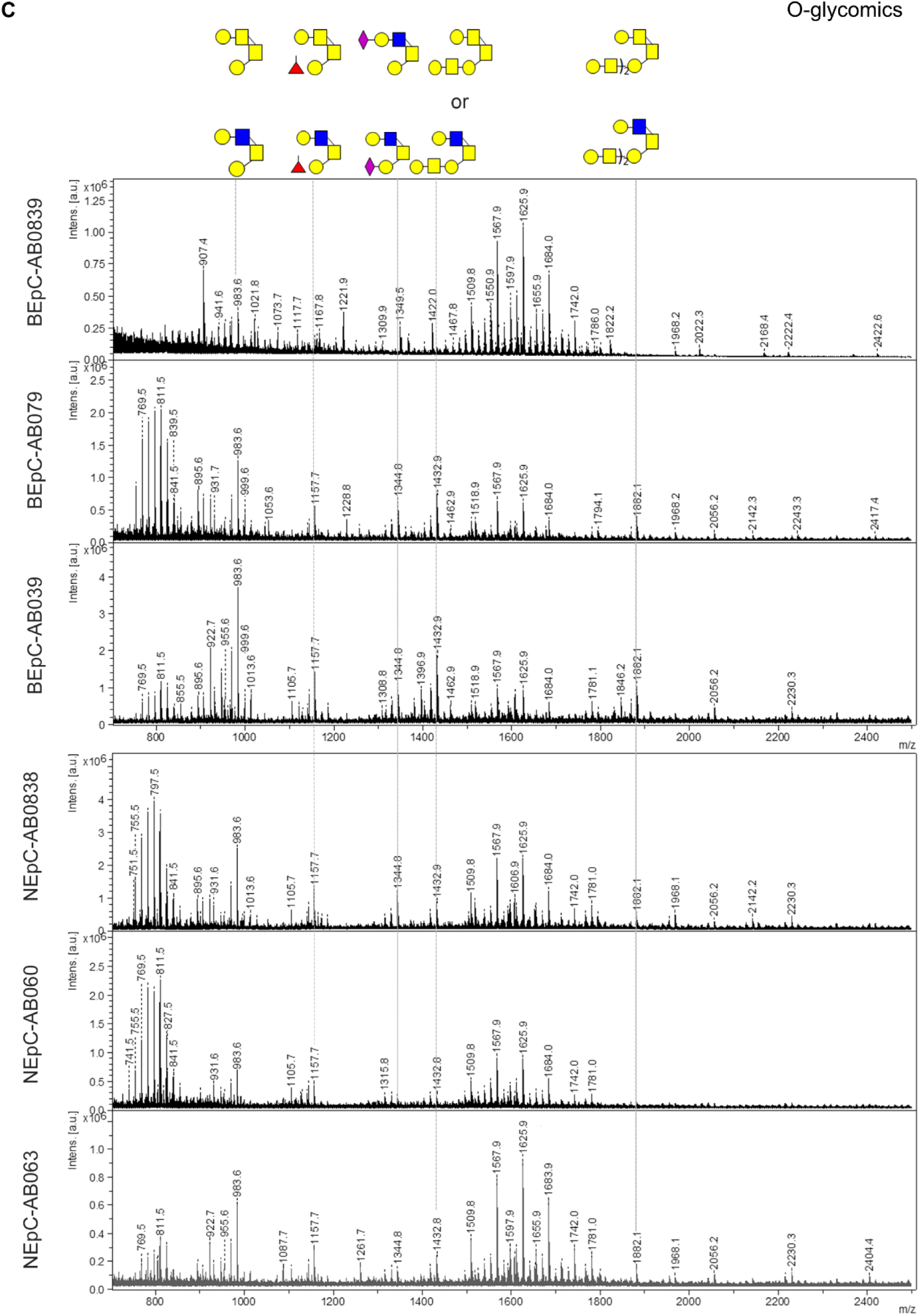
Annotated N-glycomic (A, B) and O-glycomic (C) spectra from bronchial and nasal mucus. Top to bottom: BEpC_AB051. BEpC_AB079, BEpC_AB0839, NEpC_AB038, NEpC_AB060, and NEpC_AB063. The annotation is based on the mammalian glycan biosynthesis pathway. Color codes used to describe the glycans follow the Consortium of Functional Genomics nomenclature: purple diamond: N-acetylneuraminic acid, blue square: GlcNac, green circle: mannose, yellow circle: galactose, red triangle: fucose.)_x_ indicates the number of the respective structure.

**Suppl. Figure 4.**
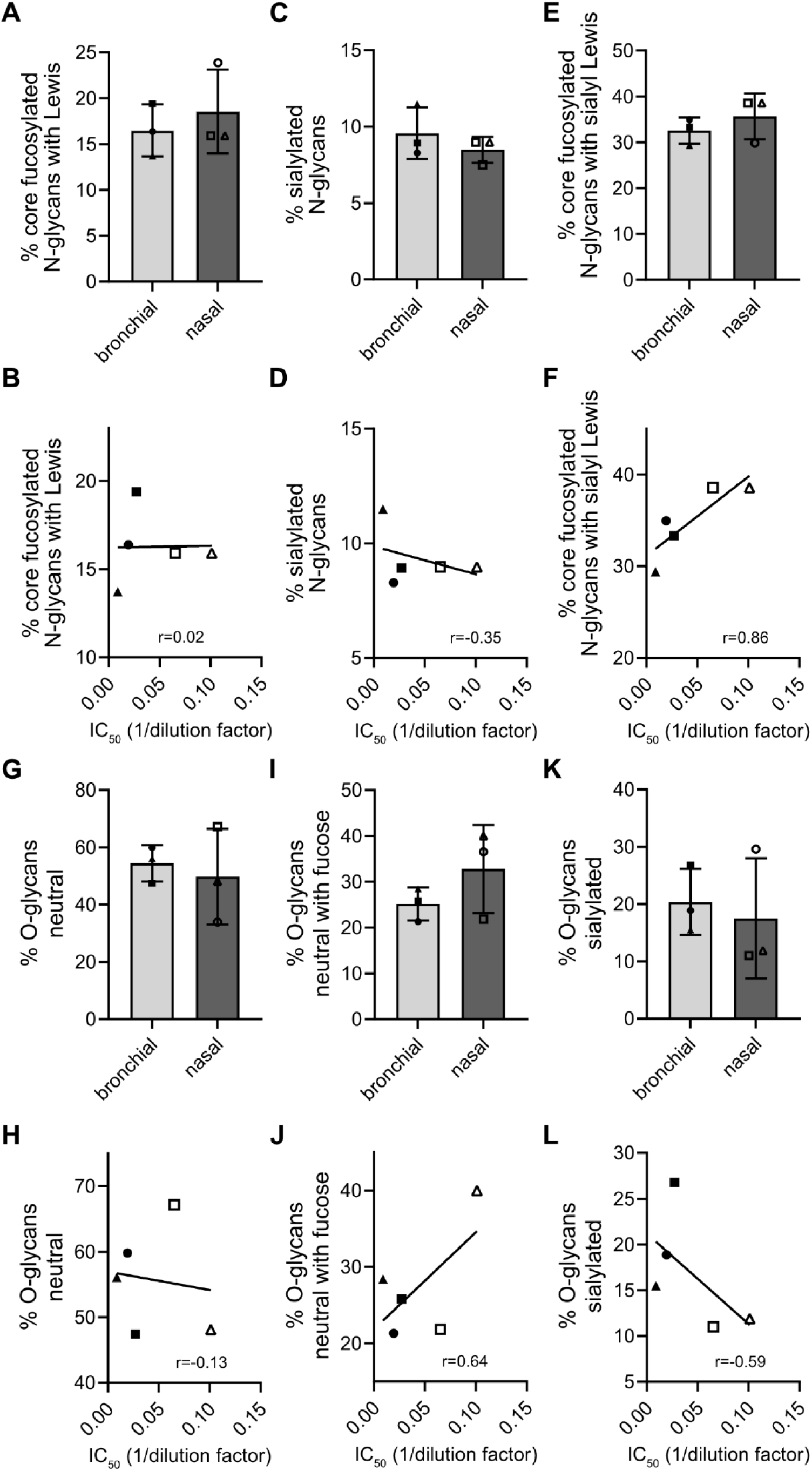
**A, C, E** Abundance of N-glycans depicted as percentage of total N-glycan abundance. Shown are N-glycans with a fucosylated core plus Lewis structure (A), sialylated glycans (B), and fucosylated N-glycans with a sialylated Lewis core (C). **B, D, F** Correlation between N-glycan subtype abundance in mucus (A, C, E) and IC_50_ of mucus against A/Brisbane/59/2007. **G, I, K** Abundance of O-glycans depicted as percentage of total O-glycan abundance. Shown are neutral O-glycans (H), neutral O-glycans with fucose (I), and sialylated O-glycans (K). **H, J, L** Correlation between O-glycan subtype abundance in mucus (G,I, K) and IC_50_ of mucus against A/Brisbane/59/2007 (Fig 2E). r describes the direction and strength of the correlation. Bronchial mucus samples are depicted as closed symbols (AB051=circle, AB079=square, AB0839=triangle); nasal mucus samples are depicted as open symbols (AB038=circle, AB060=square, AB063=triangle).

**Suppl. Table 1.**
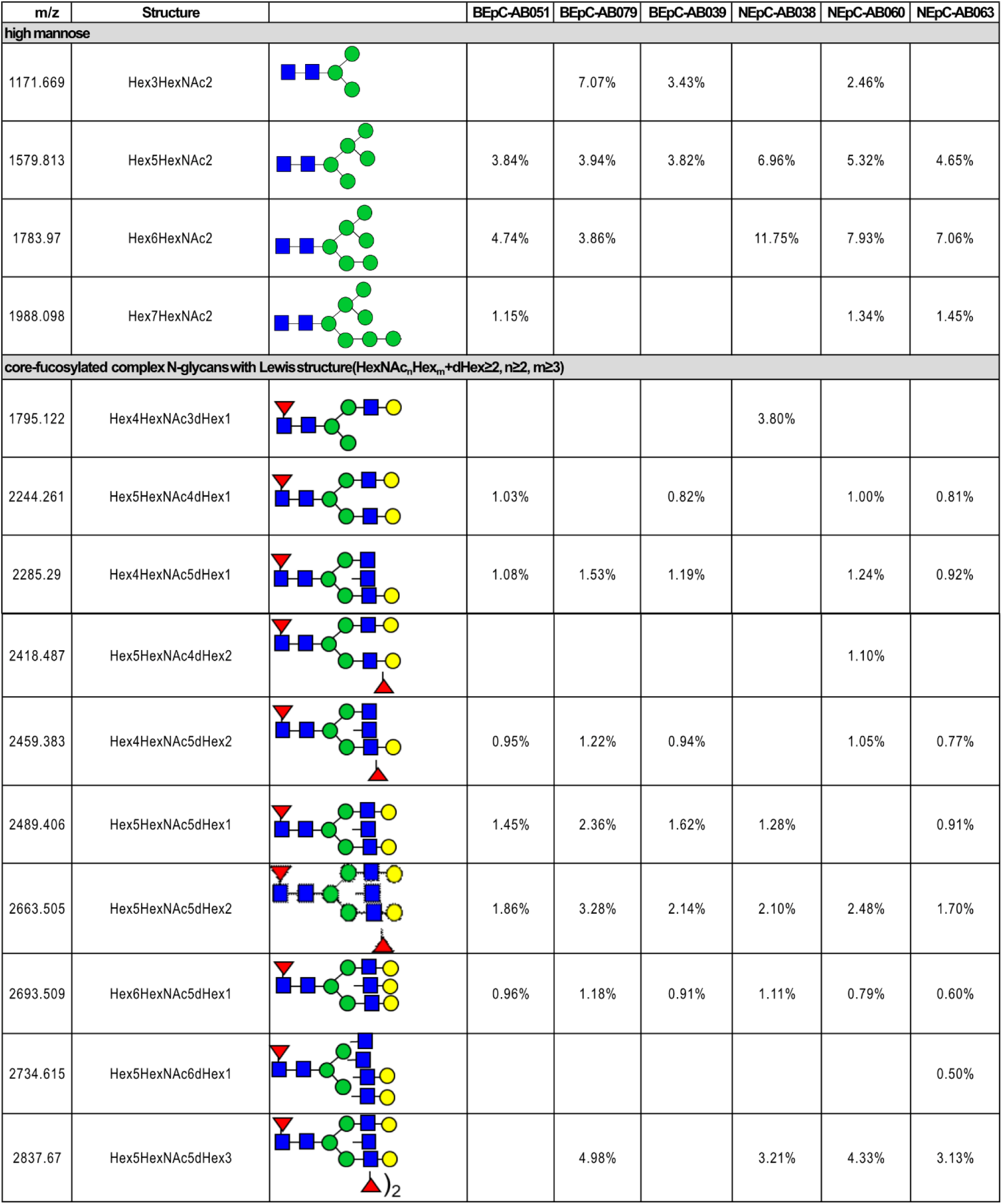
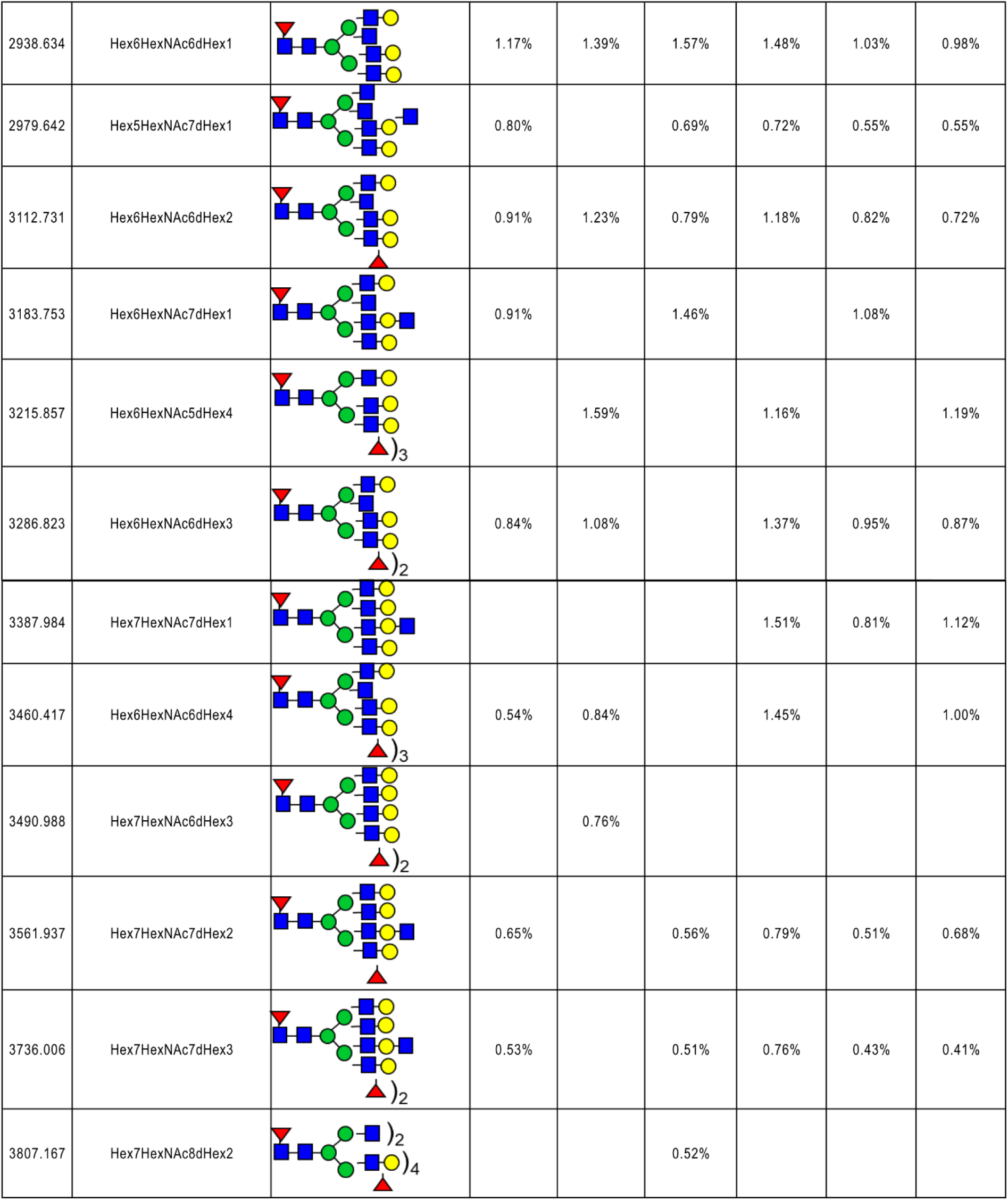
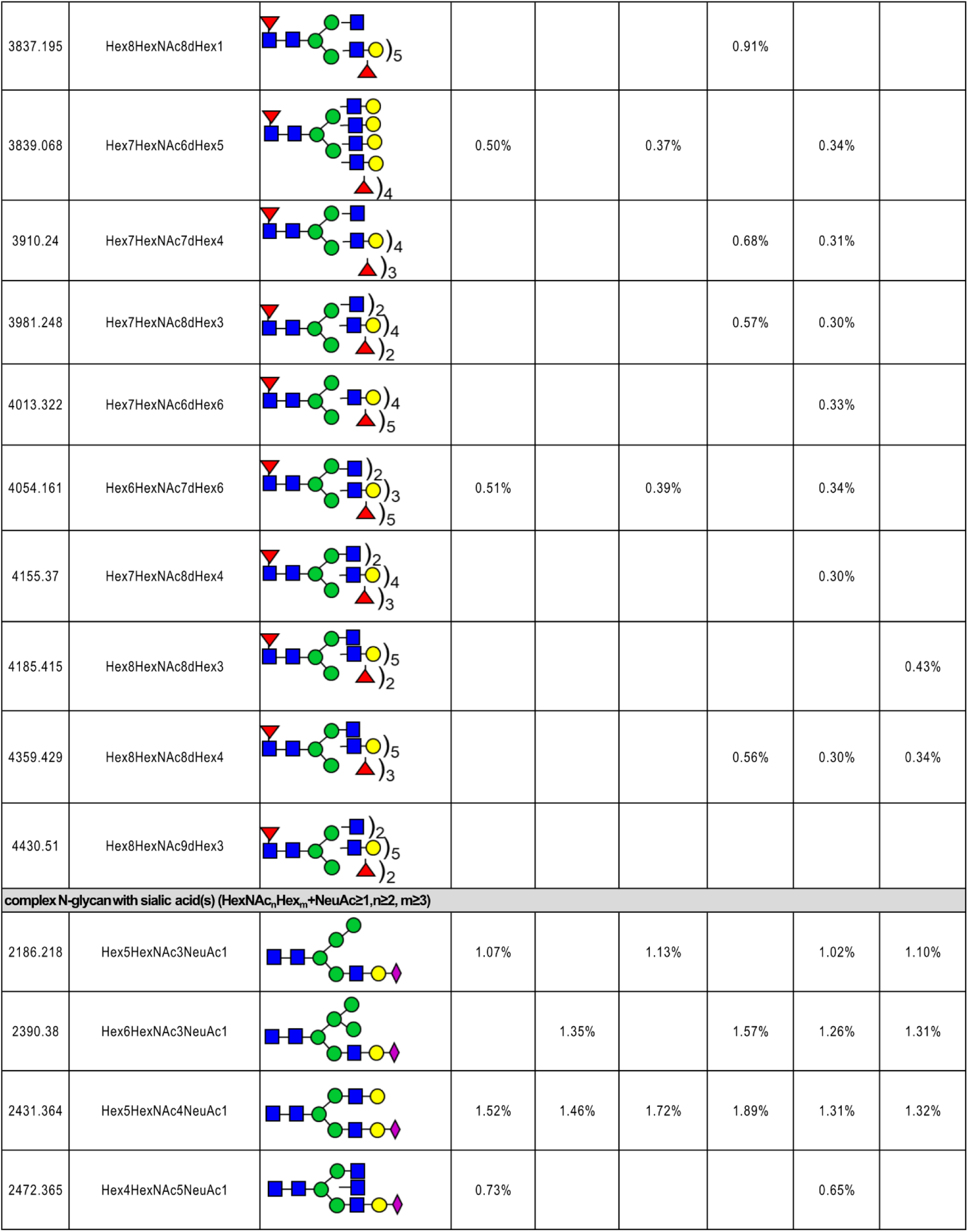
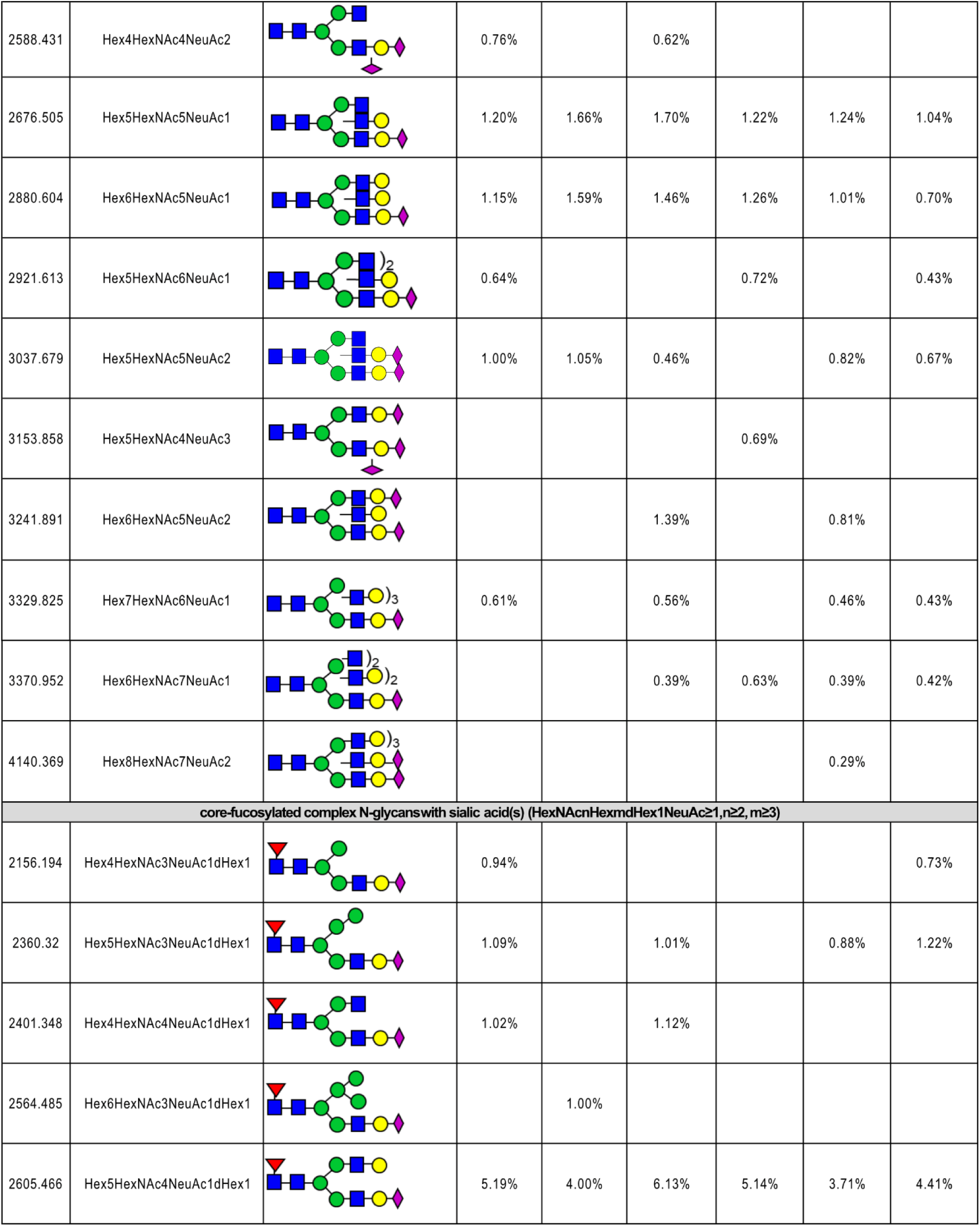
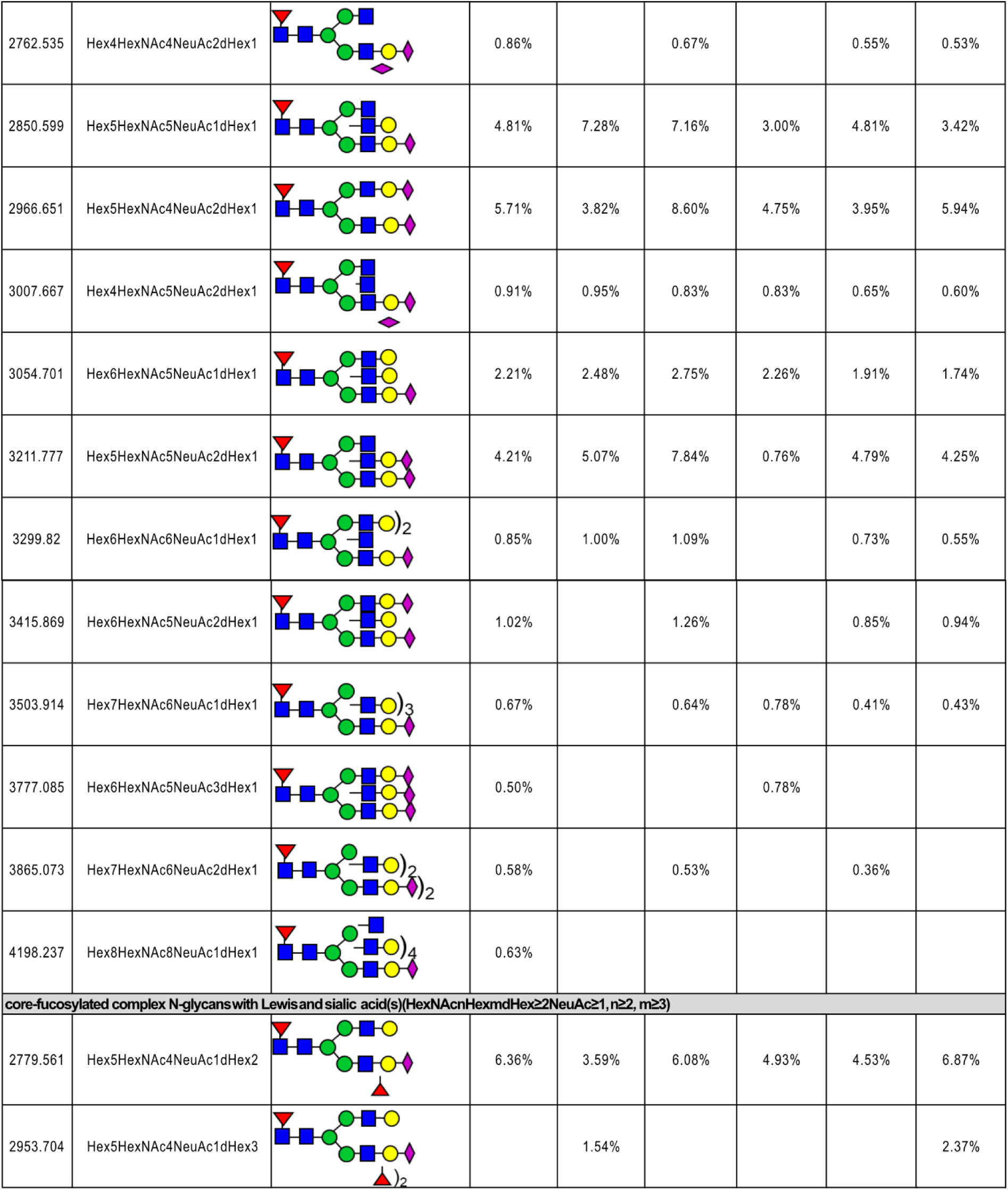
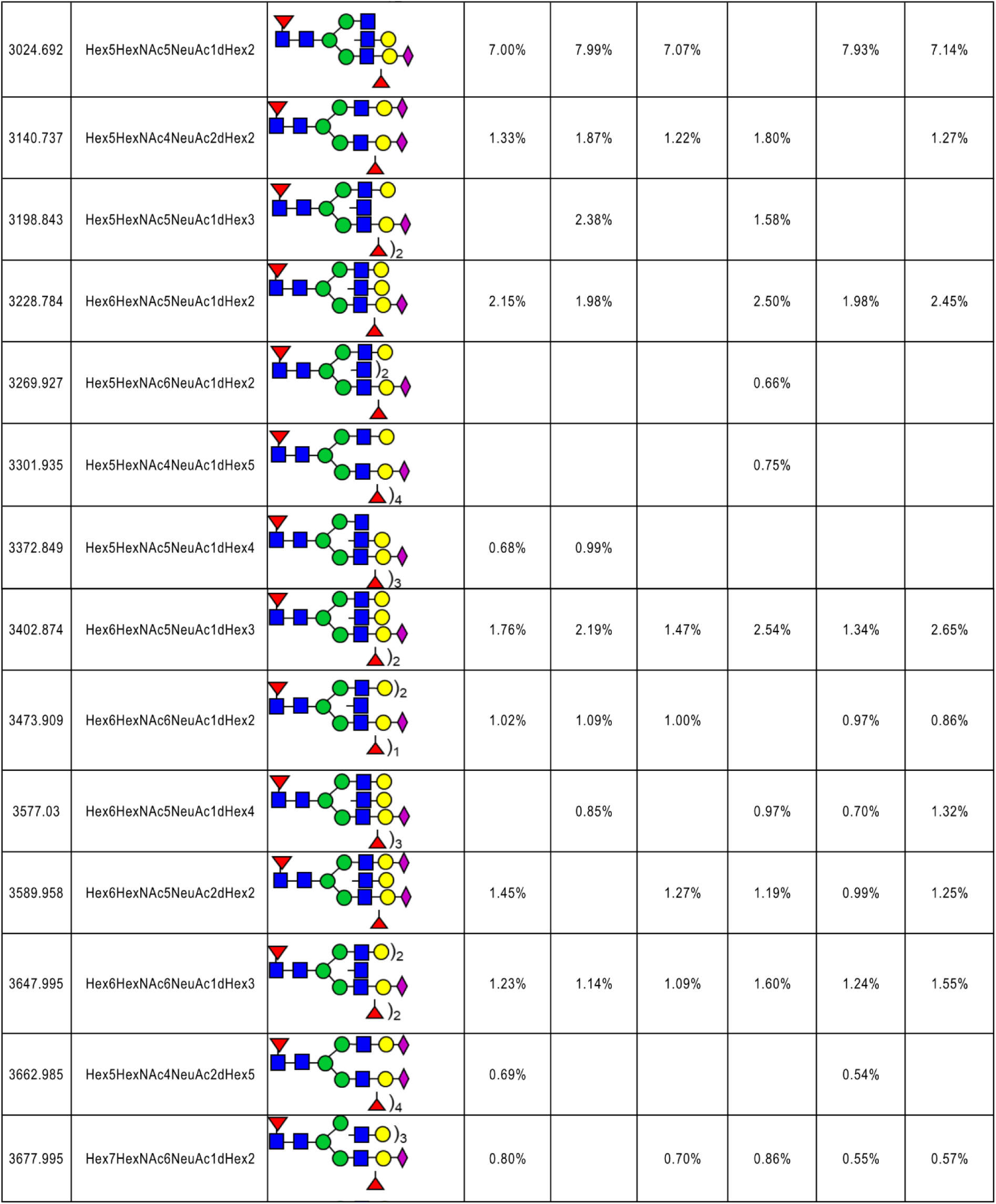
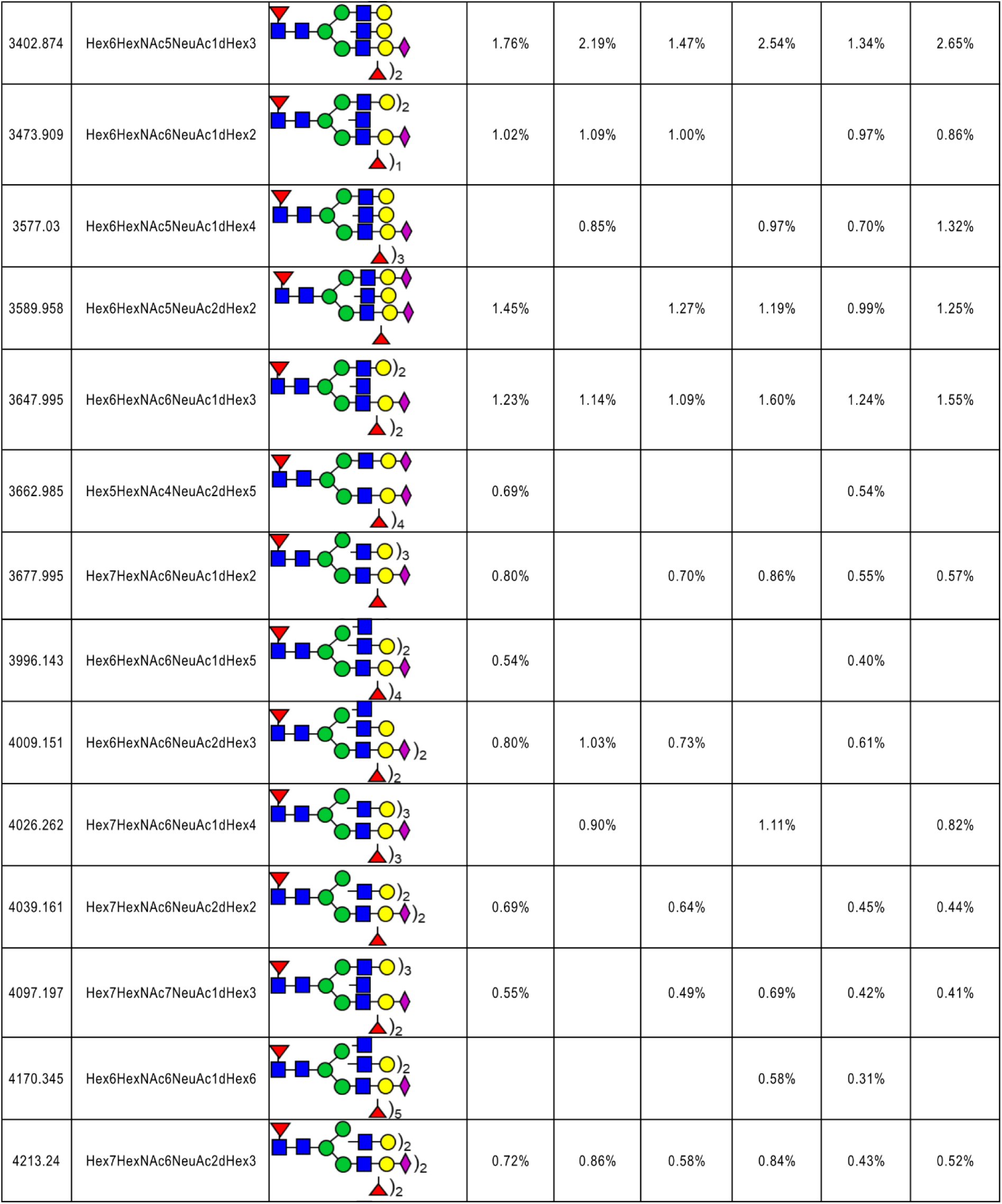
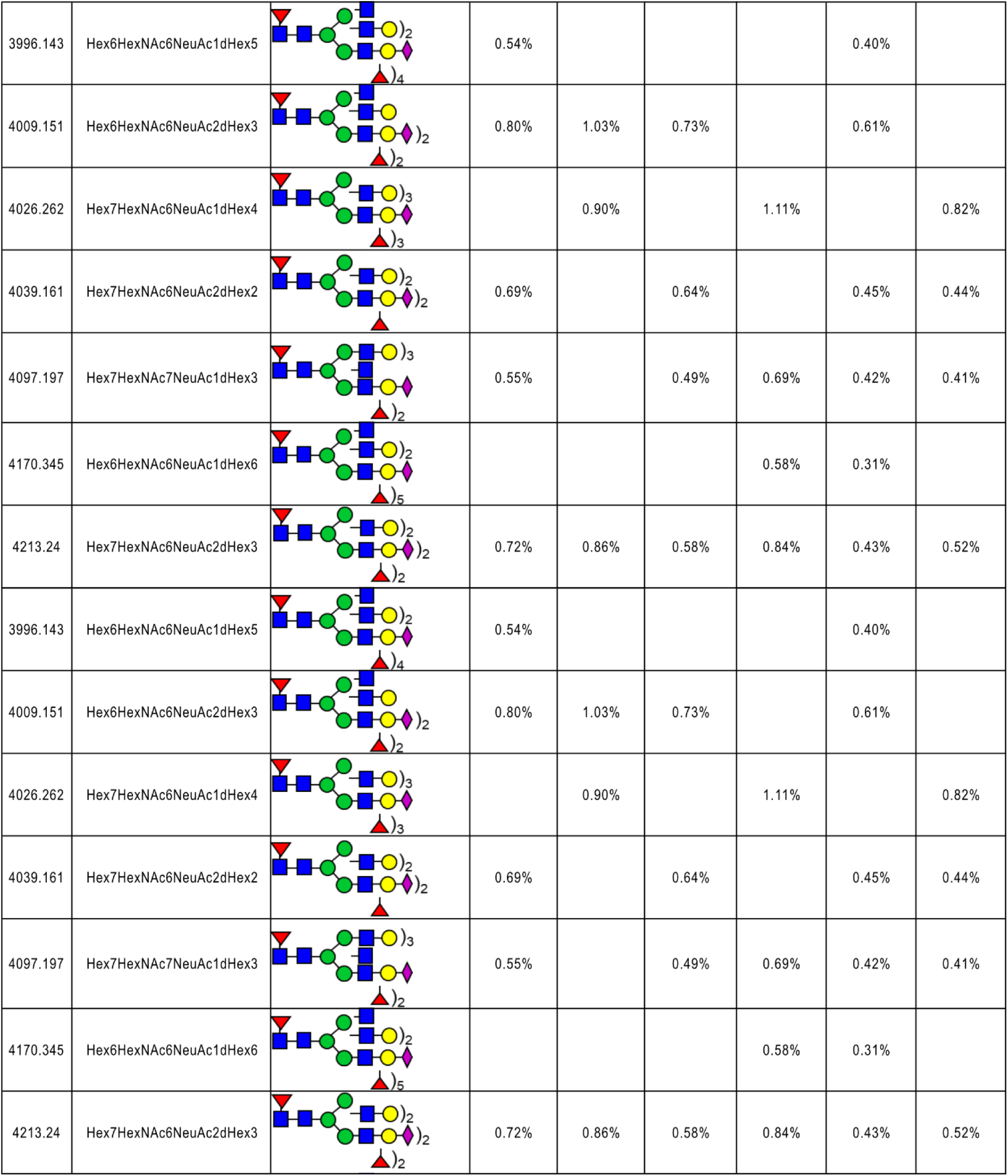
N-glycan repertoire of bronchial and nasal mucus. The summary of observed mass-to-charge ratio (m/z) and relative abundance (in percent) of N-glycans in samples. The structures were grouped into five: high mannose N-glycans, core-fucosylated complex N-glycans with Lewis structure, complex N-glycan with sialic acid(s), core-fucosylated complex N-glycans with sialic acid(s) and core-fucosylated complex N-glycans with Lewis and sialic acid(s). The principle of grouping is shown in the table. For example, core-fucosylated complex N-glycans with Lewis structure were grouped based on HexNAcnHexm+dHex≥2, n≥2, m≥3. n was the number of HexNAc and m was the number of Hexose. Since the core structure of N-glycan contains HexNAc2Hex3, n should be more than 2 and m should be more than 3. The annotation is based on the mammalian glycan biosynthesis pathway. Color codes used to describe the glycans follow the Consortium of Functional Genomics nomenclature: purple diamond: N-acetylneuraminic acid, blue square: GlcNac, green circle: mannose, yellow circle: galactose, red triangle: fucose.)_x_ indicates the number of the respective structure.

**Suppl. Table 2.** O-glycan repertoire of bronchial and nasal mucus. The summary of observed mass-to-charge ratio (m/z) and relative abundance (in percent) of O-glycans in mucus samples. The structures were grouped into three: neutral O-glycans, neutral O-glycans with fucose, and sialylated O-glycans.

| m/z | Structure | BEpC-AB051 | BEpC-AB079 | BEpC-AB039 | NEpC-AB038 | NEpC-AB060 | NEpC-AB063 |
| --- | --- | --- | --- | --- | --- | --- | --- |
| neutral O-glycans (HexNAc2Hexn) |  |  |  |  |  |  |  |
| 983.636 | HexNAc2Hex2 | 59.82% | 23.20% | 34.67% | 24.01% | 32.34% | 45.75% |
| 1228.755 | HexNAc3Hex2 |  | 5.38% |  |  |  |  |
| 1432.864 | HexNAc3Hex3 |  | 18.84% | 21.43% | 9.88% | 15.78% | 21.42% |
| neutral O-glycans with Fucose (HexNAc2HexnFuc) |  |  |  |  |  |  |  |
| 1157.718 | HexNAc2Hex2Fuc | 21.30% | 10.02% | 13.20% | 12.28% | 24.64% | 21.82% |
| 1518.903 | HexNAc2Hex2Fuc2 |  | 8.73% | 5.96% | 10.90% |  |  |
| 1606.96 | HexNAc3Hex3Fuc |  | 7.05% | 9.25% | 13.32% | 15.35% |  |
| sialylated O-glycans (HexNAc2HexnNeuAc) |  |  |  |  |  |  |  |
| 1344.806 | HexNAc2Hex2NeuAc |  | 14.98% | 10.84% | 12.94% | 11.89% | 11.01% |
| 1794.06 | HexNAc3Hex3NeuAc |  | 5.97% | 4.66% | 7.33% |  |  |
| 1968.162 | HexNAc3Hex3NeuAcFuc | 18.88% | 5.83% |  | 9.32% |  |  |

